# Stimuli reduce the dimensionality of cortical activity

**DOI:** 10.1101/026666

**Authors:** Luca Mazzucato, Alfredo Fontanini, Giancarlo La Camera

## Abstract

The activity of ensembles of simultaneously recorded neurons can be represented as a set of points in the space of firing rates. Even though the dimension of this space is equal to the ensemble size, neural activity can be effectively localized on smaller subspaces. The dimensionality of the neural space is an important determinant of the computational tasks supported by the neural activity. Here, we investigate the dimensionality of neural ensembles from the sensory cortex of alert rats during periods of ongoing (intertrial) and stimulus-evoked activity. We find that dimensionality grows linearly with ensemble size, and grows significantly faster during ongoing activity compared to evoked activity. We explain these results using a spiking network model based on a clustered architecture. The model captures the difference in growth rate between ongoing and evoked activity and predicts a characteristic scaling with ensemble size that could be tested in high-density multi-electrode recordings. Moreover, we present a simple theory that predicts the existence of an upper bound on dimensionality. This upper bound is inversely proportional to the amount of pair-wise correlations and, compared to a homogeneous network without clusters, it is larger by a factor equal to the number of clusters. The empirical estimation of such bounds depends on the number and duration of trials and is well predicted by the theory. Together, these results provide a framework to analyze neural dimensionality in alert animals, its behavior under stimulus presentation, and its theoretical dependence on ensemble size, number of clusters, and correlations in spiking network models.

## 1 Introduction

Understanding the dynamics of neural activity and how it is generated in cortical circuits is a fundamental question in Neuroscience. The spiking activity of ensembles of simultaneously recorded neurons can be represented in terms of sequences of firing rate vectors, as shown e.g. in frontal [1–3], gustatory [4, 5], motor [6], premotor and somatosensory cortex [7]. The dimension of each firing rate vector is equal to the number of ensemble neurons *N* and the collection of rate vectors across trials takes the form of a set of points in the *N*-dimensional space of firing rates. Such points may not fill the whole space, but be restricted to lie inside a lower-dimensional subspace (see e.g. [8]). Roughly, dimensionality is the minimal number of dimensions necessary to provide an accurate description of the neural dynamics. If ensemble neurons are independent of each other, neural activities at different times will scatter around in the space of firing rate, filling a large portion of the space. In this case, dimensionality will be maximal and equal to the size of the ensemble *N*. At the other extreme, if all neurons are strongly correlated, ensemble activity localizes along a line. In this case, dimensionality is minimal and equal to one. These simple examples suggest that dimensionality captures information about the structure of a cortical circuit and the functional relations among the simultaneously recorded neurons, such as their firing rates correlation computed over timescales of hundreds of milliseconds.

Different definitions of dimensionality have been introduced for different tasks and across neural systems [8–14]. Such measures of dimensionality can shed light on the underlying neural computation; for example, they can predict the onset of an error trial in a recall task [13], or can allow the comparison of classification accuracy between different brain areas (e.g., IT vs. V4) and synthetic algorithms [12]. Here, we investigate a measure of dimensionality closely related to the firing rate correlations of simultaneously recorded neurons [10]; such correlations may provide a signature of feature-based attention [15] and other top-down cognitive factors [16]. We elucidate the dependence of dimensionality on experimental parameters, such as ensemble size and interval length, and we show that it varies across experimental conditions. We address these issues by comparing recordings of ensembles of neurons from the gustatory cortex (GC) of alerts rats to a biologically plausible network model based on neural clusters with recurrent connectivity. This model captures neural activity in GC during periods of ongoing and stimulus-evoked activity, explaining how the spatiotemporal dynamics of ensemble activity is organized in sequences of metastable states and how single-neuron firing rate distributions are modulated by stimulus presentation [5]. Here, we show that the same model expounds the observed dependence of dimensionality on ensemble size and how such dependence is reduced by the presentation of a stimulus. By comparing the clustered network model with a homogeneous network without clusters, we find that the clustered network has a larger dimensionality that depends on the number of clusters and the firing rate correlations among ensemble neurons. A simple theory explains these results and allows extrapolating the scaling of dimensionality to very large ensembles. Our theory shows that recurrent networks with clustered connectivity provide a substrate for high-dimensional neural representations, which may lead to computational advantages.

## 2 Methods

### 2.1 Experimental procedures

Adult female Long Evans rats were used for this study [5, 17]. Animals received ad lib. access to food and water, unless otherwise mentioned. Movable bundles of sixteen microwires attached to a “mini-microdrive” [17, 18] were implanted in GC (AP 1.4, ML ± 5 from bregma, DV–4.5 from dura). After electrode implantation, intraoral cannulae (IOC) were inserted bilaterally [19, 20]. At the end of the surgery a positioning bolt for restraint was cemented in the acrylic cap. Rats were given at least 7 days for recovery before starting the behavioral procedures outlined below. All experimental procedures were approved by the Institutional Animal Care and Use Committee of Stony Brook University and complied with University, state, and federal regulations on the care and use of laboratory animals. More details can be found in [17]. Rats were habituated to being restrained and to receiving fluids through IOCs, and then trained to self-deliver water by pressing a lever following a 75 dB auditory cue at a frequency of 4 KHz. The interval at which lever-pressing delivered water was progressively increased to 40 ± 3 s (ITI). During training and experimental sessions additional tastants were automatically delivered at random times near the middle of the ITI, at random trials and in the absence of the anticipatory cue. Upon termination of each recording session the electrodes were lowered by at least 150 μm so that a new ensemble could be recorded. A computer-controlled, pressurized, solenoid-based system delivered ∼ 40*μ*l of fluids (opening time ∼ 40 ms) directly into the mouth through a manifold of 4 polymide tubes slid into the IOC. The following tastants were delivered: 100 mM NaCl, 100 mM sucrose, 100 mM citric acid, and 1 mM quinine HCl. Water (∼ 50μl) was delivered to rinse the mouth clean through a second IOC five seconds after the delivery of each tastant. Each tastant was delivered for at least 6 trials in each condition. Upon termination of each recording session the electrodes were lowered by at least 150μm so that a new ensemble could be recorded. Evoked activity periods were defined as the interval after tastant delivery (time *t* = 0 in our figures) and before water rinse (time *t* = 5 s). Only trials in which the tastants were automatically delivered were considered for the analysis of evoked activity, to minimize the effects of cue-related expectations [17]. Ongoing activity periods were defined as the 5 s-long intervals at the end of each inter-trial period. The behavioral state of the rat was monitored during the experiment for signs of disengagement. Erratic lever pressing, inconstant mouth movements and fluids dripping from the mouth indicated disengagement and led to the termination of the experiment. In addition, since disengagement from the task is also reflected in the emergence of high power oscillations in local field potentials, occurrences of such periods were removed offline and not analyzed further [21].

### 2.2 Data analysis

Single neuron action potentials were amplified, bandpass filtered (at 300 – 8 KHz), digitized and recorded to a computer (Plexon, Dallas, TX). Single units of at least 3: 1 signal-to-noise ratio were isolated using a template algorithm, cluster cutting techniques and examination of inter-spike interval plots (Offline Sorter, Plexon, Dallas, TX). All data analyses and model simulations were performed using custom software written in Matlab (Mathworks, Natick, MA, USA), Mathematica (Wolfram Research, Champaign, IL), and C. Starting from a pool of 299 single neurons in 37 sessions, neurons with peak firing rate lower than 1 Hz (defined as silent) were excluded from further analysis, as well as neurons with a large peak around the 6 – 10 Hz in the spike power spectrum, which were considered somatosensory [17, 22, 23]. Only ensembles with 3 or more simultaneously recorded neurons were further analyzed (167 non-silent, non-somatosensory neurons from 27 ensembles). We analyzed ongoing activity in the 5 seconds interval preceding either the auditory cue or taste delivery, and evoked activity in the 5 seconds interval following taste delivery in trials without anticipatory cue, wherein significant taste-related information is present [24].

### 2.3 Hidden Markov Model (HMM) analysis

Here we briefly outline the procedure used in [5], see this reference and [4, 7, 25] for further details. Under the HMM, a system of N recorded neurons is assumed to be in one of a predetermined number of hidden (or latent) states [26, 26]. Each state *m* is defined as a vector of *N* firing rates *v_i_(m), i* = 1,…,*N*, one for each simultaneously recorded neuron. In each state, the neurons were assumed to discharge as stationary Poisson processes (Poisson-HMM). We matched the model to the data segmented in 1-ms bins (see below). In such short bins, we found that typically at most one spike was emitted across all simultaneously recorded neurons. If more than one neuron fired an action potential in a given bin, only one (randomly chosen) was kept for further analysis (this only occurred in a handful of bins per trial) [25]. We denote by *y_i_(t)* the spiking activity of the *i*-th neuron in the interval [*t, t + dt*], *y_i_(t)* = 1 if the neuron emitted a spike and *y_i_(t)* = 0 otherwise. Denoting with *S_t_* the hidden state of the ensemble at time *t*, the probability of having a spikes from neuron *i* in a given state *m* in the interval [*t, t + dt*] is given by *p(y_i_(t)* = 1|*S_t_ = m*) = 1 – *e^v_i_(m)dt^*.

The firing rates *v_i_(m)* completely define the states and are also called “emission probabilities” in HMM parlance. The emission and transition probabilities were found by maximization of the log-likelihood of the data given the model via the expectation-maximization (EM), or Baum-Welch, algorithm [26], a procedure known as “training the HMM”. For each session and type of activity (ongoing vs. evoked), ensemble spiking activity from all trials was binned at 1 ms intervals prior to training assuming a fixed number of hidden states *M* [4, 25]. For each given number of states *M*, the Baum-Welch algorithm was run 5 times, each time with random initial conditions for the transition and emission probabilities. The range of hidden states *M* for the HMM analyses were *M_min_* = 10 and *M_max_* = 20 for spontaneous activity, and *M_min_* = 10 and *M_max_* = 40 for evoked activity. Such numbers were based on extensive exploration of the parameter space and previous studies [4, 5, 7, 25, 27]. For evoked activity, each HMM was trained on all four tastes simultaneously. Of the models thus obtained, the one with largest total likelihood *M*\* was taken as the best HMM match to the data, and then used to estimate the probability of the states given the model and the observations in each bin of each trial (a procedure known as “decoding”). During decoding, only those hidden states with probability exceeding 80% in at least 50 consecutive bins were retained (henceforth denoted simply as “states”). State durations were approximately exponentially distributed with median duration 0.60 s (95% CIs: 0.07 – 4.70) during ongoing activity and 0.30 s (0.06 – 2.80) during evoked activity [5]. The firing rate fits *v_i_(m)* in each trial were obtained from the analytical solution of the maximization step of the Baum-Welch algorithm,

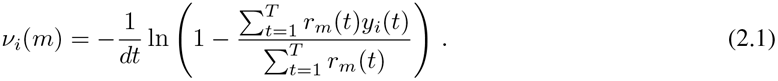

Here, [*y_i_*(1),…, *y_i_(T)*] is the spike train of the *i*-th neuron in the current trial, and *T* is the total duration of the trial. *r_m_(t)* = *P(S_t_ = m|y*(1),…, *y(T)*) is the probability that the hidden state *S_t_* at time *t* is *m*, given the observations.

### 2.4 Dimensionality measure

We defined the dimensionality of the neural activity as [10]

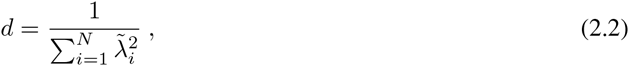

where the 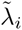 are the principal eigenvalues expressed as fractions of the total amount of variance explained, i.e. 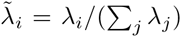, where *λ_j_* are the eigenvalues of the covariance matrix of the firing rates (see below). The dimensionality can be computed exactly in some relevant special cases. The calculation is simplified by the observation that Eq. (2.2) is equivalent to

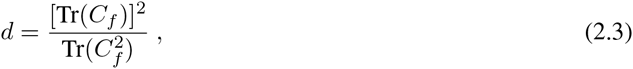

where *C_f_* is the true covariance matrix of the firing rate vectors, 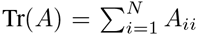 is the trace of matrix *A*, and 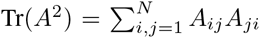. We consider in the following only the case of firing rates in equal bins, hence we can replace *C_f_* with the covariance matrix of the spike counts *C* in the definition of *d*:

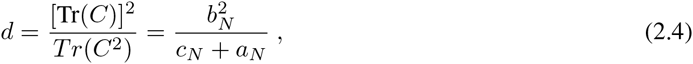

where for later convenience we have introduced the notation

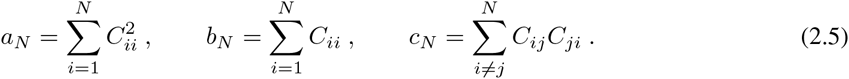

Note that *d* does not depend on the distribution of firing rates, but only on their covariance, up to a common scaling factor.

*Dimensionality in the case of uniform pair-wise correlations*. When all the pair-wise correlations *r_ij_* are identical, *r_ij_ = ρ* for all *i ≠ j*,

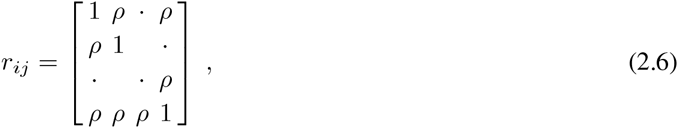

we have 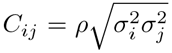 for *i ≠ j*, where 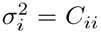 is the spike count variance. In this case, we find from Eq. (2.5) that

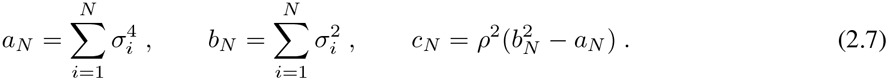

and the dimensionality, Eq. (2.4), is given by

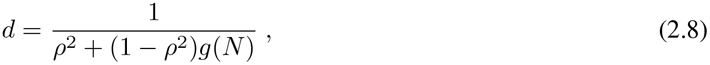

where

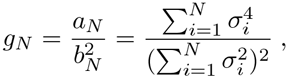

Note that since both *a_N_* and *b_N_* scale as *N* when *N* is large, in general 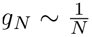 for large *N*. If all spike counts have equal variance, σ_*i*_ = σ, we find exactly 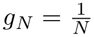:

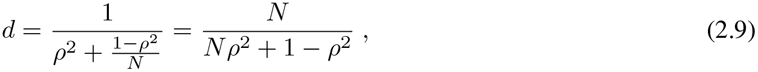

and the dependence of *d* on the variance drops out. Note that for uncorrelated spike counts (*ρ* = 0) this formula gives *d = N*, whereas for any finite correlation we find the upper bound 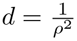. For *N* > 1, the dimensionality is inversely related to the amount of pair-wise correlation *ρ*.

Consider the case where spike counts have variances 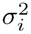 drawn from a probability distribution with mean 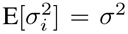 and variance 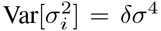, and the pair-wise correlation coefficients *r_ij_*, for *i ≠ j*, are drawn from a distribution with mean 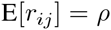 and variance Var[*r_ij_*] = *δρ*^2^. In such a case one can evaluate Eq. (2.4) approximately by its Taylor expansion around the mean values of the quantities in Eqs. (2.5). At leading order in *N* one

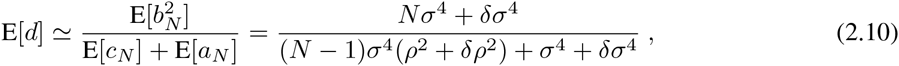

where E[.] denotes expectation. To obtain this result we have used the definitions in Eq. (2.5), from which

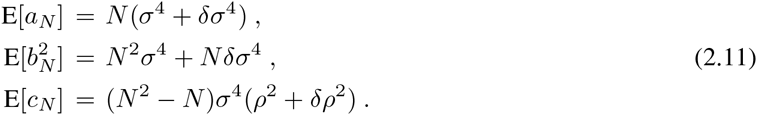

and the fact that, given a random vector *X_i_* with mean *μ_i_* and covariance *C_ij_*, and a constant symmetric matrix *A_ij_*, the expectation value of the quadratic form Σ*_i,j_ X_i_A_ij_ X_j_* is

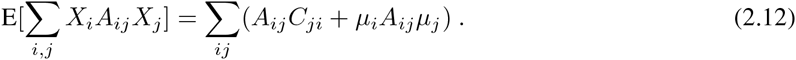

In the case of uncorrelated spike counts (*ρ* = 0, *δρ* = 0), dimensionality still depends linearly on the ensemble size *N*, but with a smaller slope *σ*^4^/(*σ*^4^ + *δσ*^4^) < 1 compared to the case of equal variances (Eq. (2.9) with *ρ* = 0).

*Dimensionality in the case of neural clusters*. Given an ensemble of *N* neurons arranged in *Q* clusters (motivated by the model network described later in Sec. 2.8), we created ensembles of uncorrelated spike trains for *N ≤ Q* and correlated within each cluster for *N > Q*. Thus, if *N ≤ Q* the correlation matrix is the *N* x *N* identity matrix. If *N > Q*, the (*Q* + 1)-th neuron was added to the first cluster, with correlation *ρ* with the other neuron of the cluster, and uncorrelated to the neurons in the remaining clusters. The (*Q* + 2)-th neuron was added to the second cluster, with correlation *ρ* with the other neuron of the second cluster, and uncorrelated to the neurons in the remaining clusters, and so on. Similarly, the (*2Q* + *p*)-th neuron (*p ≤ Q)* was added to the *p*-th cluster, with pair-wise correlation *ρ* with the other neurons of the same cluster, but no correlation with the neurons in the remaining clusters; and so on. In general, for *N* = *mQ* + *p* neurons (where *m* = [*N/Q*]_ ≥ 1 is the largest integer smaller than *N/Q*), the procedure picked *m* + 1 neurons per cluster for the first *p* cluster and *m* neurons per cluster for the remaining *Q – p* clusters, with uniform pair-wise correlations *ρ* in the same cluster while neurons from different clusters were uncorrelated. The resulting correlation matrix *r* was block diagonal

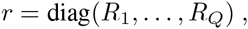

where each of the *Q* blocks contains the correlations of neurons from the same cluster. Inside each block *R_i_*, the off-diagonal terms are equal to the uniform within-cluster correlation *ρ:*

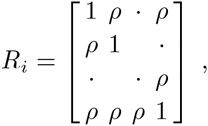

The first *p* blocks have size (*m* + 1) x (*m* + 1) and the last *Q – p* blocks have size *m x m*, so that (*m* + 1)*p* + *m(Q – p) = N*. The remaining elements of matrix *r* (representing pair-wise correlations of neurons belonging to different clusters) were all zero. Recalling that *C_ij_ = *r*_ij_σ_i_σ_j_*, one finds Tr(*C*) = *pb*_*m*+1_ + (*Q – p*)*b_m_* and 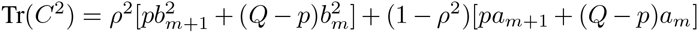, where *a_n_* and *b_n_* are defined in Eq. (2.7), from which one obtains

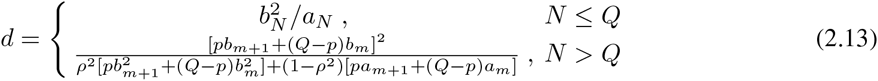

In the approximation where all neurons have the same variance this simplifies to

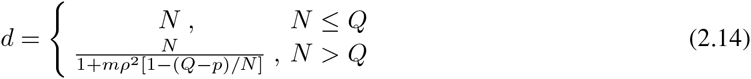

Recall that in the formulae above *m* and *p* depend on *N*. For finite *ρ*, Eq. (2.14) predicts the bound *d ≤ Q/ρ*^2^ for any *N*, with this value reached asymptotically for large *N*. When single neuron variances 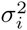 are drawn from a distribution with mean 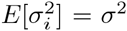 and variance 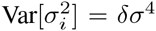, an expression for the dimensionality can be easily obtained from Eq. (2.13) at leading order around the expectation values in Eq. 2.5 (not shown), with a procedure similar to that use to obtain Eq. (2.10).

### 2.5 Pair-wise correlations

Given neuron *i* and neuron *j*’s spike trains, we computed the spike count correlation coefficient *r_ij_*,

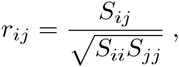

where *S* is the sample covariance matrix of the spike counts estimated as

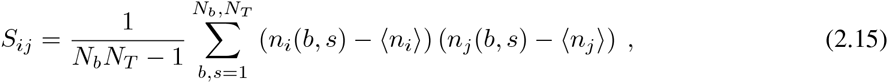

where *n_i_ (b, s)* is the spike count of neuron *i* in bin *b* and trial *s*. The sum goes over all *N_b_* bins and over all *N_T_* trials in a session, whereas 〉ni〈 is the average across trials and bins for neuron *i*. In the main text and figures we present results obtained with a bin size of 200 ms, but have performed the same analyses with bin sizes varying from 10 ms to 5 seconds (see Results for details).

Significance of the correlation was estimated as follows [28]: *N_shuffle_* = 200 trial-shuffled correlation coefficients 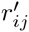 were computed, then a *p*-value was determined as the fraction of shuffled coefficients 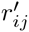 whose absolute value exceeded the absolute value of the experimental correlation, 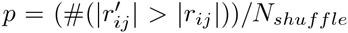. For example, a correlation *r* was significant at *p* = 0.05 confidence level if no more than 10 shuffled correlation coefficients out of 200 exceeded *r*. The pair-wise correlations of firing rates vectors computed in bins of fixed duration *T* were given by Eq. (2.15) with *n_i_(b, s)* replaced by *n_i_(b, s)/T*. Instead, correlations of firing rates vectors inside hidden states (which have variable duration) were estimated after replacing *n_i_(b, s)* in Eq. (2.15) with *v_i_(m, s)*, the firing rate of neuron *i* in state m in trial *s*. For each trial *s*, this quantity was computed according to Eq. (2.1).

### 2.6 Estimation of dimensionality

The eigenvalues *λ_j_* in Eq. (2.2) were found with a standard Principal Component Analysis (PCA) of the set of all firing rate vectors [29]. The firing rate vectors were obtained via the HMM analysis (see Eq. (2.1)); all data from either ongoing or evoked activity were used. For the analysis of Fig. 3E, where the duration and number of trials were varied, only the firing rate vectors of the HMM states present in the given trial snippet were used (even if present for only a few ms). When firing rate vectors in hidden states were not available (mainly, in “shuffled” datasets and in asynchronous homogeneous networks, see below for details), the firing rates were computed as spike counts in *T* = 200 ms bins divided by *T, n_i_(b, s)/T*, where *n_i_(b, s)* is as defined in Eq. (2.15) (Fig. 3F, 3G, 6E, 7D and 9A). Dimensionality values were averaged across 20 simulated sessions for each ensemble size *N*; in each session, 40 trials of 5 s duration, resulting in *N_T_* = 1,000 bins, were used (using bin widths of 50 to 500 ms did not change the results). Note that for the purpose of computing the dimensionality (Eq. (3)), it is equivalent to use either the binned firing rate *n_i_(b, s)/T* or the spike count *n_i_(b, s)*.

In our data, *d* roughly corresponded to the number of principal components explaining between 80 to 90% of the variance. However, note that all eigenvalues are retained in our definition of dimensionality given in Eq. (2.2) above.

*Shuffled datasets*. The dimensionality of the data as a function of ensemble size *N* was validated against surrogate datasets constructed by shuffling neurons across different sessions while matching the empirical distribution of ensemble sizes. Comparison analyses between empirical and shuffled ensembles were trial-matched using the minimal number of trials per condition across ensembles, and then tested for significant difference with the Mann-Whitney test on samples obtained from 20 bootstrapped ensembles. Neurons whose firing rate variance exceeded the population average by two standard deviations were excluded (8/167 of non-silent, non-somatosensory neurons).

*Dependence on the number of trials: simulations (Fig. 7E, 8A)*. The estimate of d from data depends on the number and duration of the trials (Fig. 3E and Eq. (2.17) below). To investigate this phenomenon in a simple numerical setting we generated *N* x *N_T_* “nominal” firing rates, thought of as originating from *N* neurons, each sampled *N_T_* times (trials). The single firing rates were sampled according to a log-normal distribution with equal means and covariance leading to Eq. (2.8), i.e., 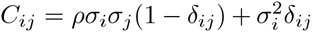, with *δ_ij_* = 1 if *i = j*, and zero otherwise (note that the actual distribution used is immaterial since the dimensionality only depends on the covariance matrix, see Eq. (2.4)). We considered the two cases of equal variance for all ensemble neurons, *σ_i_ = σ* for all *i* (Fig. 8A) or variances *σ_i_* sampled from a log-normal distribution (Fig. 8A and “+” in Fig. 7E). The same *N* and *N_T_* as used for the analysis of the model simulations in Fig. 7D were used (where the “trials” were *N_T_* bins of 200 ms in 40 intervals of 5 second duration for each ensemble size *N*). The covariance of the data thus generated was estimated according to Eq. (2.15), based on which the dimensionality Eq. (2.4) was computed. The estimated dimensionality depends on *N* and *N_T_* and was averaged across 100 values of *d*, each obtained as explained above. Note that in this simplified setting increasing the duration of each trial is equivalent to adding more trials, i.e., the effect of having a trial 400 ms long producing 2 firing rates (one for each 200 ms bin) is equivalent to having two trials of 200 ms duration. In the general case, the effect of trial duration on *d* will depend on how trial duration affects the variance and correlations of the firing rates.

*Dependence on the number of trials: theory*. The dependence of dimensionality on the number of trials can be computed analytically under the assumption that *N* ensemble neurons generate spike counts *n_i_*, for *i* = 1,…, *N*, distributed according to a multivariate Gaussian. Since we are interested in the spike-count covariance Eq. (2.15), we can assume the spike-count distribution to have zero mean and true covariance *C_ij_*. The matrix *M* = (*N_T_* – 1) • *S^(N_T_)^*, where *S^(N_T_)^* is the covariance matrix Eq. (2.15) sampled from *N_T_* trials, is distributed according to a Wishart distribution *W_N_*(*C_ij_, N_T_* – 1) with *N_T_* – 1 degrees of freedom [30]. Since the variance of the Wishart distribution,

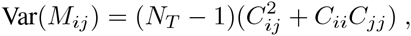

is proportional to *N_T_*, we obtain the variance of the entries of the sample covariance as

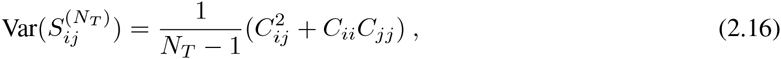

to be used in the estimator of *d* (from Eq. (2.4))

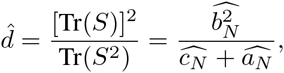

where 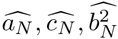 are given by Eq. (2.5) with *C* replaced by *S*. With a calculation similar to that used to obtain Eq. (2.10), to leading order in *N* and *N_T_* one finds

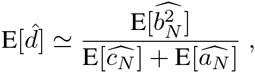

with

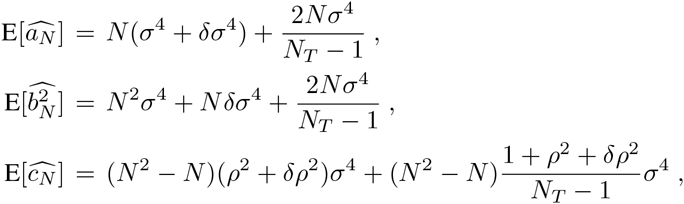

where we also used Eqs. (2.11) and (2.12), with 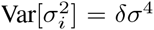 and Var[*r_ij_*] = *δρ*^2^, for *i* ≠ *j*. In conclusion, one finds

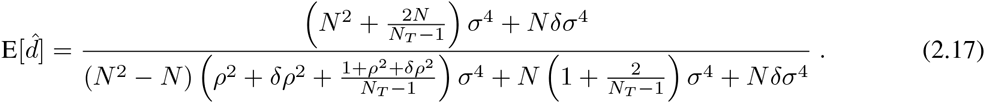

*Model fitting*. The dependence of the datas dimensionality on ensemble size *N* was fitted by a straight line via standard least-squares,

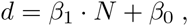

separately for ongoing and evoked activity (Fig. 3B-D and 6B-D). Comparison between the dimensionality of evoked and ongoing activity was carried out with a 2-way ANOVA with condition (evoked vs. ongoing) and ensemble size (*N*) as factors. Since *d* depends on the number and duration of the trials used to estimate the covariance matrix (Fig. 3E and Eq. (2.17)), we matched both the number of trials and trial length in comparisons of ongoing and evoked dimensionality. If multiple tastes were used, the evoked trials were each matched to a random subset of an equal number of ongoing trials.

The dependence of dimensionality d on ensemble size *N* in a surrogate dataset of Poisson spike trains with mean pairwise correlation *ρ* (generated according to the algorithm described in the next section) was modeled as Eq. (2.17) with *δρ*^2^ = *αρ*^2^ and *δσ*^4^ = *σ*^4^ = *β* (Fig. 7D, dashed lines); *N_T_* was fixed to 1000 (40 trials of 5 seconds each, segmented in 200 ms bins). The parameters *α, β* were tuned to fit all Poisson trains simultaneously on datasets with *N* = 5,10,…, 100 and *ρ* = 0,0.01,0.05,0.1,0.2, with 20 ensembles for each value (Fig. 7D; only the fits for *ρ* = 0,0.1,0.2 are shown). A standard non-linear least-squares procedure was used [31].

### 2.7 Generation of correlated Poisson spike trains

Ensembles of independent and correlated Poisson spike trains were generated for the analysis of Fig. (7). Ensembles of independent stationary Poisson spike trains with given firing rates *ν_i_* were generated by producing their interspike intervals according to an exponential distribution with parameter *ν_i_*. Stationary Poisson spike trains with fixed pairwise correlations (but no temporal correlations) were generated according to the method reported in [32], that we briefly outline below. We split each trial into 1 ms bins and consider the associated binary random variable *X_i_(t)* = 1 if the *i*-th neuron emitted a spike in the *t*-th bin, and *X_i_(t)* = 0 if no spike was emitted. These samples were obtained by first drawing a sample from an auxiliary *N*-dimensional Gaussian random variable *U ∼ N*(γ, Λ) and then thresholding it into 0 and 1: *X_i_* = 1 if *U_i_* > 0, and *X_i_* = 0 otherwise. Here, γ = {γ_1_,γ_2_,…, γ_*N*_} is the mean vector and Λ = {Λ_*ij*_} is the covariance matrix of the *N*-dimensional Gaussian variable *U*. For appropriately chosen parameters γ_*i*_ and Λ_*ij*_ the method generates correlated spike trains with the desired firing rates *ν_i_* and pairwise spike count correlation coefficients *r_ij_*.

The prescription for *γ_i_* and Λ*_ij_* is most easily expressed as a function of the desired probabilities *μ_i_* of having a spike in a bin of width *dt, μ_i_* = *P*(*X_i_(t)* = 1), and the pairwise covariance *S_ij_* of the random binary vectors *X_i_(t)* and *X_j_(t)*, from which *γ_i_* and Λ*_ij_* can be obtained by inverting the following relationships:

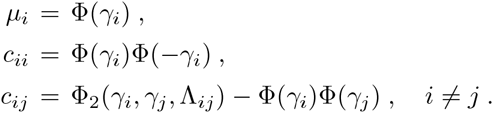

Here, Φ(*x*) is the cumulative distribution of a univariate Gaussian with mean 0 and variance 1 evaluated at *x*, and Φ_2_ (*x, y*, Λ) is the cumulative distribution of a bivariate Gaussian with means 0, variances 1 and covariance Λ evaluated at (*x, y*) (note that the distributions Φ and Φ_2_ are unrelated to the *N*-dimensional Gaussian *U* ∼ *N*(*γ*, Λ)). Without loss of generality we imposed unit variances for *U_i_*, i.e. Λ_*ii*_ = 1.

We related the spike probabilities *μ_i_* to the firing rates *ν_i_* as *μ_i_* = 1 – *e-^ν_i_dt^*, with 1 – *μ_i_* being the probability of no spikes in the same bin. When *dt* approaches zero, *μ_i_* ≃ *ν_i_dt* and the spike trains generated as vectors of binary random variables by sampling *U ∼ N*(γ, Λ) will approximate Poisson spike trains (*dt* = 1 ms bins were used). In order to have a fair comparison with the data generated by the spiking network model (described in the next section), the mean firing rates of the Poisson spike trains were matched to the average firing rates obtained from the simulated data. Since γ and Λ were the same in all bins, values of *X_i_(t)* and *X_i_(s)* were independent for *t ≠ s* (i.e., the spike trains had no temporal correlations). As a consequence, the random binary vectors have the same pair-wise correlations as the spike counts, and the *c_ij_* are related to the desired 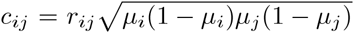, where *μ_i_*(1 – *μ_i_*) is the variance of *X_i_*. See [32] for further details.

### 2.8 Spiking network model

We modeled the data with a recurrent spiking network of *N* = 5000 randomly connected leaky integrate-and-fire (LIF) neurons, of which 4000 excitatory (*E*) and 1000 inhibitory (*I*). Connection probability *p_βα_* from neurons in population *α* ∈ *E, I* to neurons in population *β ∈ E, I* were *p_EE_* = 0.2 and *p_EI_ = p_IE_ = p_ii_* = 0.5; a fraction *f* = 0.9 of excitatory neurons were arranged into *Q* different clusters, with the remaining neurons belonging to an unstructured (“background”) population [33]. Synaptic weights *J_βα_* from neurons in population *α ∈ E, I* to neurons in population *β ∈ E, I* scaled with *N* as 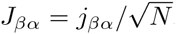, with *j_βα_* constants having the following values (units of mV): *j_EI_* = 3.18, *j_IE_* = 1.06, *j_ii_* = 4.24, *j_EE_* = 1.77. Within an excitatory cluster synaptic weights were potentiated, i.e. they took average values of 〉J〈_+_ = *J_+_ j_EE_* with *J*_+_ > 1, while synaptic weights between units belonging to different clusters were depressed to average values 〉J〈_ = *J*_*j_EE_*, with *J*_ − = 1 – *γf*(*J*_+_ – 1) < 1, with γ = 0.5. The latter relationship between *J*_+_ and *J*_-_helps to maintain balance between overall potentiation and depression in the network [33].

Below spike threshold, the membrane potential V of each LIF neuron evolved according to

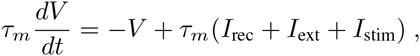

with a membrane time constant *τ_m_* = 20 ms for excitatory and 10 ms for inhibitory units. The input current was the sum of a recurrent input *I_rec_*, an external current *I_ext_* representing an ongoing afferent input from other areas, and an external stimulus *I_stim_* representing e.g. a delivered taste during evoked activity only. In our units, a membrane capacitance of 1 nF is set to 1. A spike was said to be emitted when *V* crossed a threshold *V_thr_*, after which *V* was reset to a potential *V_reset_* = 0 for a refractory period of *τ_ref_* = 5 ms. Spike thresholds were chosen so that, in the unstructured network (i.e., with *J*_+_ = *J*_ − = 1), the *E* and *I* populations had average firing rates of 3 and 5 spikes/s, respectively [33]. The recurrent synaptic input 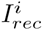 to unit *i* evolved according to the dynamical equation

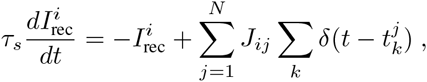

where 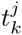 was the arrival time of *k*-th spike from the *j*-th pre-synaptic unit, and *τ_s_* was the synaptic time constant (3 and 2 ms for *E* and I units, respectively), resulting in an exponential post-synaptic current in response to a single spike, 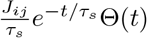, where Θ(*t*) = 1 for *t* ≥ 0, and Θ(*t*) = 0 otherwise. The ongoing external current to a neuron in population *α* was constant and given by

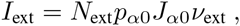

where 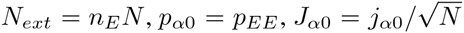 with *j_E0_* = 0.3, *j*_*I*0_ = 0.1, and *ν_ext_* = 7 spikes/s. During evoked activity, stimulus-selective units received an additional input representing one of the four incoming stimuli. The stimuli targeted combinations of neurons as observed in the data. Specifically, the fractions of neurons responsive to *n* = 1, 2, 3 or all 4 stimuli were 17%(27/162), 22%(36/162), 26%(42/162), and 35%(57/162) [5, 24]. Each stimulus had constant amplitude *ν_stim_* ranging from 0 to 0.5*ν_ext_*. In the following we measure the stimulus amplitude as percentage of *ν_ext_* (e.g., “10%” corresponds to *ν_stim_* = 0.1*ν_ext_*). The onset of each stimulus was always *t* = 0, the time of taste delivery. The stimulus current to a unit in population *α* was constant and given by

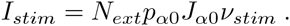

### 2.9 Mean field analysis of the model

The stationary states of the spiking network model in the limit of large *N* were found with a mean field analysis [5, 33–36]. Under typical conditions, each neuron of the network receives a large number of small post-synaptic currents (PSCs) per integration time constant. In such a case, the dynamics of the network can be analyzed under the diffusion approximation within the population density approach. The network has *α* = 1,…, *Q* + 2 sub-populations, where the first *Q* indices label the *Q* excitatory clusters, *α = Q* + 1 labels the “background” units, and *α = Q* + 2 labels the homogeneous inhibitory population. In the diffusion approximation [37–39], the input to each neuron is completely characterized by the infinitesimal mean *μ_α_* and variance 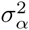 of the post-synaptic potential (see [5] for the expressions of the infinitesimal mean and variance for all subpopulations).

Parameters were chosen so that the network with *J*_+_ = *J*_-_ = 1 (where all *E → E* synaptic weights are equal) would operate in the balanced asynchronous regime [28, 40, 41], where incoming contributions from excitatory and inhibitory inputs balance out, neurons fire irregular spike trains with weak pair-wise correlations.

The unstructured network has only one dynamical state, i.e., a stationary point of activity where all *E* and I neurons have constant firing rate *ν_E_* and *ν_I_*, respectively. In the structured network (where *J*_+_ > 1), the network undergoes continuous transitions among a repertoire of states, as shown in the main text. To avoid confusion between network activity states and HMM states, we refer to the former as network “configurations” instead of states. Admissible networks configurations must satisfy the *Q*+2 self-consistent mean field equations [33]

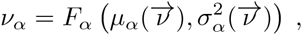

where 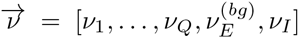 is the firing rate vector and 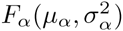 is the current-to-rate response function of the LIF neurons. For fast synaptic times, i.e. 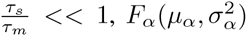 is well approximated by [42, 43]

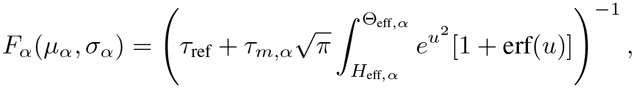

where

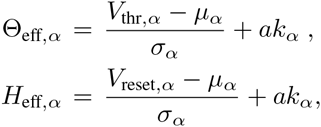

where 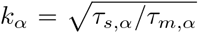 is the square root of the ratio of synaptic time constant to membrane time constant, and 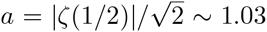. This theoretical response function has been fitted successfully to the firing rate of neocortical neurons in the presence of in vivo-like fluctuations [44–47].

The fixed points 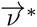 of the mean field equations were found with Newtons method [48]. The fixed points can be either stable (attractors) or unstable depending on the eigenvalues *λ_α_* of the stability matrix

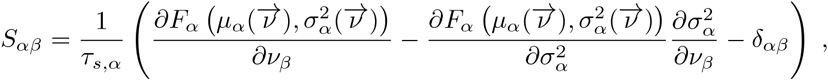

evaluated at the fixed point 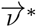 [49]. If all eigenvalues have negative real part, the fixed point is stable (attractor). If at least one eigenvalue has positive real part, the fixed point is unstable. Stability is meant with respect to an approximate linearized dynamics of the mean and variance of the input current:

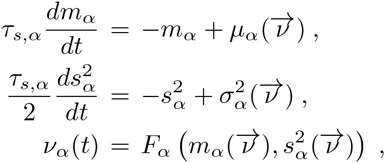

where *μ_α_* and 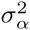 are the stationary values for fixed 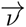 given earlier. For fast synaptic dynamics in the asynchronous balanced regime, these rate dynamics are in very good agreement with simulations ([50] see [51, 52] for more detailed discussions).

### 2.10 Metastable configurations in the network model

The stable configurations of a network with an infinite number of neurons were obtained in the mean field approximation of the previous section and are shown in Fig. 4B for *Q* = 30 and a range of values of the relative potentiation parameter *J*_+_. Above the critical point *J*_+_ = 4.2, stable configurations characterized by a finite number of active clusters emerge (grey lines; the number of active clusters is reported next to each line). For a given *J*_+_, the firing rate is the same in all active clusters and is inversely proportional to the total number of active clusters. Stable patterns of firing rates are also found in the inhibitory population (red lines), in the inactive clusters (having low firing rates; grey dashed lines), and in the unstructured excitatory population (dashed blue lines). For a fixed value of *J*_+_, multiple stable configurations coexist with different numbers of active clusters. For example, for *J*_+_ = 5.3, stable configurations with up to 7 active clusters are stable, each configuration with different firing rates. This generates multistable firing rates in single neurons, i.e., the property, also observed in the data, that single neurons can attain more than 2 firing rates across states [5]. Note that if *J*_+_ ≤ 5.15 an alternative stable configuration of the network with all clusters inactive (firing rates < 10 spikes/s) is also possible (single brown line). Strictly speaking, the configurations in Fig. 4B are stable only in a network containing an infinite number of uncorrelated neurons. In a finite network (or when neurons are strongly correlated) these configurations can lose stability due to strong fluctuations which ignite transitions among the different configurations. Full details are reported in [5].

### 2.11 Model simulations and analysis of simulated data

The dynamical equations of the LIF neurons were integrated with the Euler algorithm with a time step of *dt* = 0.1 ms. We simulated 20 different networks (referred to as “sessions” in the following) during both ongoing and evoked activity. We chose four different stimuli per session during evoked activity (to mimic taste delivery). Trials were 5 seconds long. The HMM analyses for Figs. 2 and 5 were performed on ensembles of randomly selected excitatory neurons with the same procedure used for the data (see previous section “Hidden Markov Model (HMM) analysis”). The ensemble sizes were chosen so as to match the empirical ensemble sizes (3 to 9 randomly selected neurons). For the analysis of Fig. 8B, random ensembles of increasing size (from 5 to 100 neurons) were used from simulations with *Q* = 30 clusters. When the ensemble size was less than the number of clusters (*N ≤ Q*), each neuron was selected randomly from a different cluster; when ensemble size was larger than the number of clusters, one neuron was added to each cluster until all clusters were represented, and so on until all *N* neurons had been chosen. To allow comparison with surrogate Poisson spike trains, the dimensionality of the simulated data was computed from the firing rate vectors in *T* = 200 ms bins as explained in Sec. 2.4. For control, the dimensionality was also computed from the firing rate vectors in hidden states obtained from an HMM analysis, obtaining qualitatively similar results.

## 3 Results

### 3.1 Dimensionality of the neural activity

We investigate the dimensionality of sequences of firing rate vectors generated in the GC of alert rats during periods of ongoing or evoked activity (see Section 2.1). To provide an intuitive picture of the meaning of dimensionality adopted in this paper, consider the firing rate vectors from *N* simultaneously recorded neurons. These vectors can occupy, a priori, the entire *N*-dimensional vector space minimally required to describe the population activity of *N* independent neurons. However, the sequence of firing rate vectors generated by the neural dynamics may occupy a subspace that is spanned by a smaller number *m < N* of coordinate axes. For example, the data obtained by the ensemble of three simulated spike counts in Fig. 1 mostly lie on a 2D space, the plane shaded in gray. Although 3 coordinates are still required to specify all data points, a reduced representation of the data, such as that obtained from PCA, would quantify the dimension of the relevant subspace as being close to 2. To quantify this fact we use the following definition of dimensionality [10]

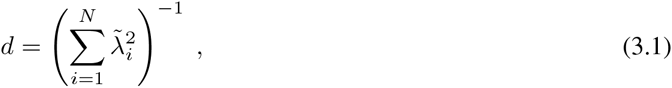

where *N* is the ensemble size and 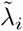 are the normalized eigenvalues of the covariance matrix, each expressing the fraction of the variance explained by the corresponding principal component (see Section 2.4 for details). According to this formula, if the first n eigenvalues express each a fraction 1/*n* of the variance while the remaining eigenvalues vanish, the dimensionality is *d = n*. In less symmetric situations, *d* reflects roughly the dimension of the linear subspace explaining most variance about all data points. In the example of the data on the gray plane of Fig. 1, *d* = 1.8, which is close to 2, as expected. Similarly, data points lying mostly along the blue and red straight lines in Fig. 1 have a dimensionality of 0.9, close to 1. In all cases, *d* > 0 and *d ≤ N*, where *N* is the ensemble size.

The blue and red data points in Fig. 1 were obtained from a fictitious scenario where neuron 1 and neuron 2 were selective to surrogate stimuli A and B, respectively, and are meant to mimic two possible evoked responses. The subspace containing responses to both stimuli A and B would have a dimensionality *d_A+B_* = 1.7, similar to the dimensionality of the data points distributed on the grey plane (meant instead to represent spike counts during ongoing activity in the same fictitious scenario). Thus, a dimensionality close to 2 could originate from different patterns of activity, such as occupying a plane or two straight lines. Other and more complex scenarios are, of course, possible. In general, the dimensionality will reflect existing functional relationships among ensemble neurons (such as pair-wise correlations) as well as the response properties of the same neurons to external stimuli. The pictorial example of Fig. 1 caricatures a stimulus-induced reduction of dimensionality, as found in the activity of simultaneously recorded neurons from the GC of alert rats, as we show next.

**Figure 1.**
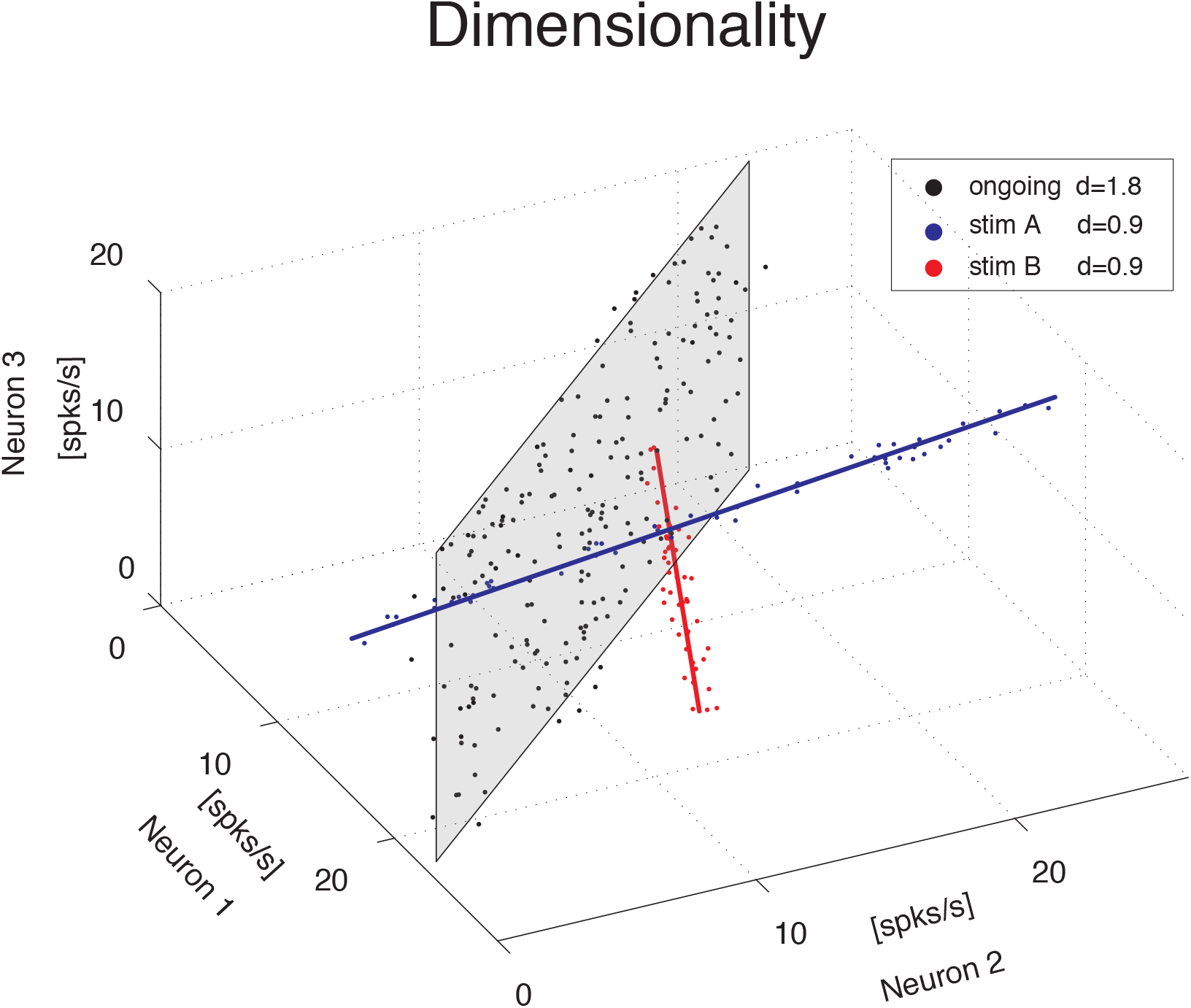
Dimensionality of the neural representation. Pictorial representation of the firing rate activity of an ensemble of *N* = 3 neurons. Each dot represents a three-dimensional vector of ensemble firing rates in one trial. Ensemble ongoing activity localizes around a plane (black dots cloud surrounding the shaded black plane), yielding a dimensionality of *d* = 1.8. Activity evoked by each of two different stimuli localizes around a line (red and blue dots clouds and lines), yielding a dimensionality of *d* = 0.9 in both cases.

### 3.2 Dimensionality is proportional to ensemble size

We computed the dimensionality of the neural activity of ensembles of 3 to 9 simultaneously recorded neurons in the gustatory cortex of alert rats during the 5 s inter-trial period preceding (ongoing activity) and following (evoked activity) the delivery of a taste stimulus (said to occur at time *t* = 0; see Methods). Ensemble activity in single trials during both ongoing (Fig. 2A) and evoked activity (Fig. 2B) could be characterized in terms of sequences of metastable states, where each state is defined as a collection of firing rates across simultaneously recorded neurons [4, 5]. Transitions between consecutive states were detected via a Hidden Markov Model (HMM) analysis, which provides the probability that the network is in a certain state at every 1 ms bin (Fig. 2, color-coded lines superimposed to raster plots). The ensemble of spike trains was considered to be in a given state if the posterior probability of being in that state exceeded 80% in at least 50 consecutive 1-ms bins (Fig. 2, color-coded shaded areas). Transitions among states were triggered by the co-modulation of a variable number of ensemble neurons and occurred at seemingly random times [5]. For this reason, the dimensionality of the neural activity was computed based on the firing rate vectors in each HMM state (one firing rate vector per state per trial; see Methods for details). The average dimensionality of ongoing activity across sessions was *d_ongoing_* = 2.6 ± 1.2 (mean±SD; range: [1.2, 5.0]; 27 sessions). An example of the eigenvalues for a representative ensemble of eight neurons is shown in Fig. 3A, where *d* = 4.42. The dimensionality of ongoing activity was approximately linearly related to ensemble size (Fig. 3B, linear regression, *r* = 0.4, slope *b_ongoing_* = 0.26 ± 0.12, *p* = 0.04). During evoked activity dimensionality did not differ across stimuli (oneway ANOVA, no significant difference across tastants, *p* > 0.8), hence all evoked data points were combined for further analysis. An example of the eigenvalue distribution of the ensemble in Fig. 2B is shown in Fig. 3C, where *d_evoked_* = 1.3 ∼ 1.7 across 4 different taste stimuli. Across all sessions, dimensionality was overall smaller (*d_evoked_* = 2.0±0.6, mean±SD, range: [1.1, 3.9]) and had a reduced slope as a function of *N* compared to ongoing activity (Fig. 3D, linear regression, *r* = 0.39, slope *b_evoked_* = 0.13 ± 0.03, *p* < 10^−4^). However, since dimensionality depends on the number and duration of the trials used for its estimation (Fig. 3E), a proper comparison requires matching trial number and duration for each data point, as described next.

**Figure 2.**
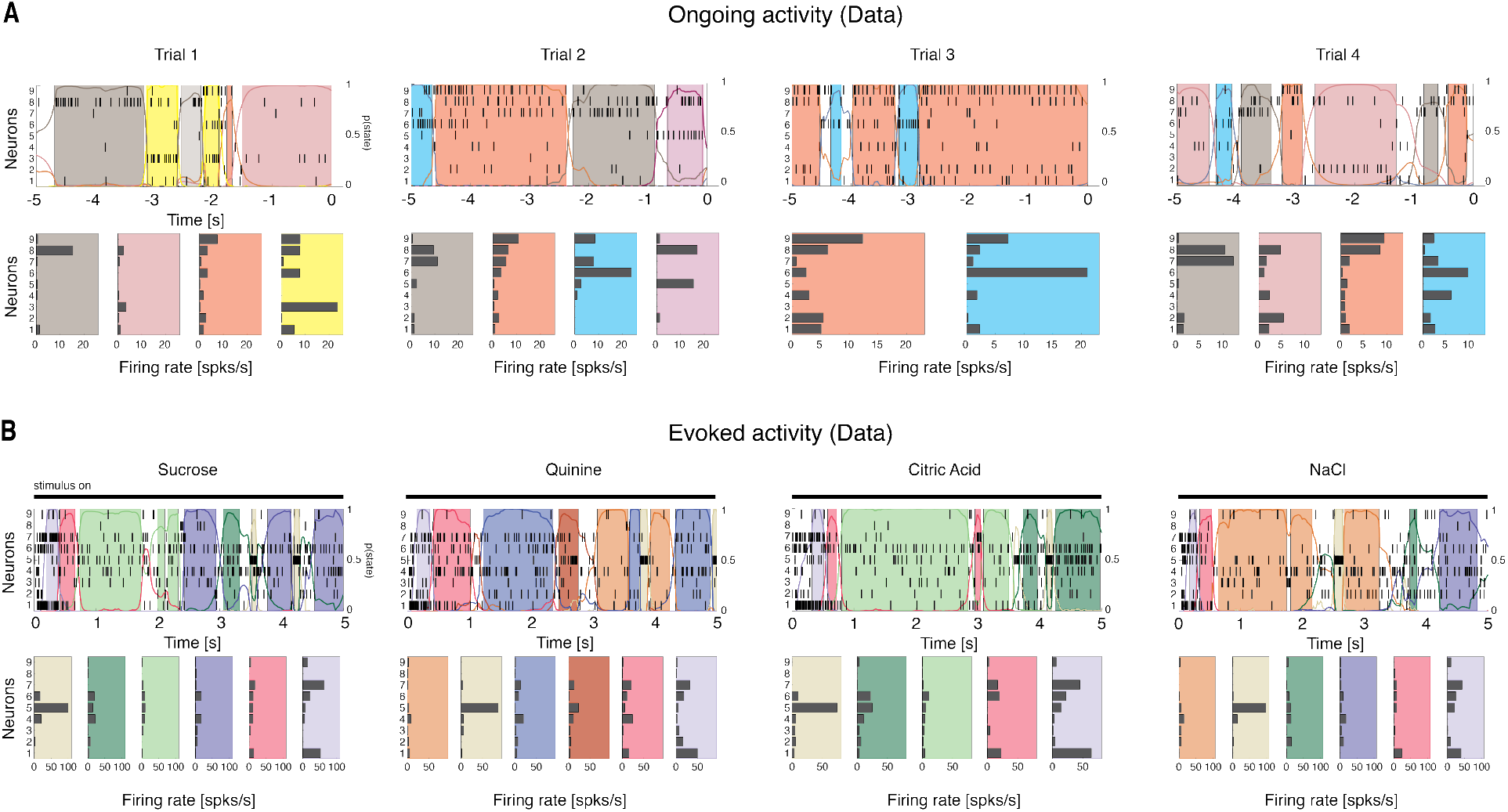
Ensemble neural activity is characterized by sequences of states. A: Upper panels: Representative trials from one ensemble of nine simultaneously recorded neurons during ongoing activity, segmented according to their ensemble states (HMM analysis, thin black vertical lines are action potentials; states are color-coded; smooth colored lines represent the probability for each state; shaded colored areas indicate intervals where the probability of a state exceeds 80%). Lower panels: Average firing rates across simultaneously recorded neurons (states are color-coded as in the upper panels). In total, 6 hidden states were found in this example session. X-axis for population rasters: time preceding the next event at (0 = stimulus delivery); Y-axis for population rasters: left, ensemble neuron index, right, probability of HMM states; X-axis for average firing rates panels: firing rates (spks/s); Y-axis for firing rate panels: ensemble neuron index. B: Ensemble rasters and firing rates during evoked activity for four different tastes delivered at *t* = 0 (the black line on top of the raster plot represents the “stimulus-on” period): sucrose, sodium chloride, citric acid and quinine (notations as in panel A). In total, eight hidden states were found in this session during evoked activity.

**Figure 3.**
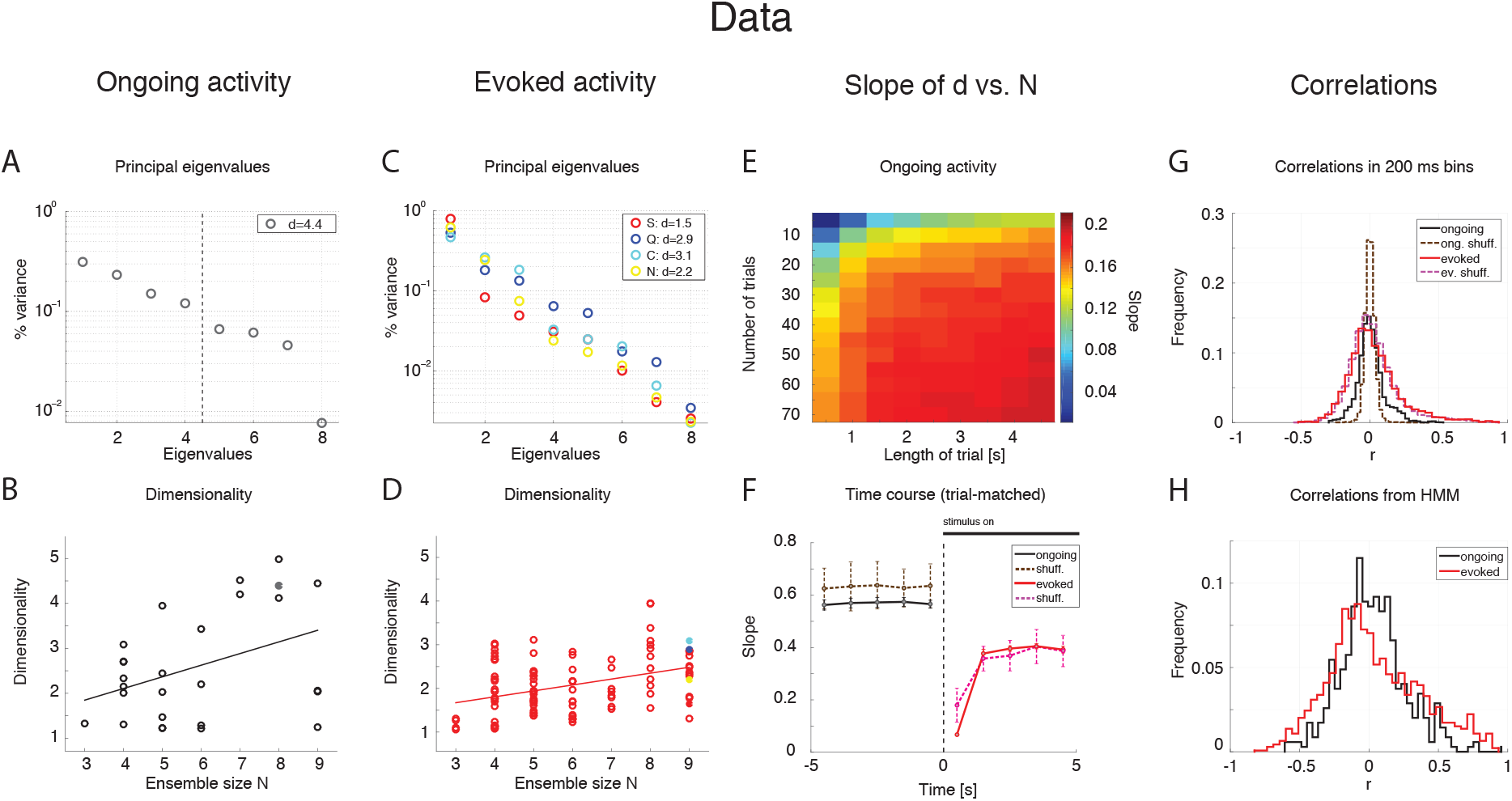
Dependence of dimensionality on ensemble size (data). A: Fraction of variance explained by each principal eigenvalue for an ensemble of 8 neurons during ongoing activity (corresponding to the filled dot in panel B) in the empirical dataset. The dashed vertical line represents the value of the dimensionality for this ensemble (*d* = 4.4). X-axis: eigenvalue number; Y-axis: fraction of variance explained by each eigenvalue. B: Dimensionality of neural activity across all ensembles in the empirical dataset during ongoing activity (circles, linear regression fit, *d = b·N+a, b* = 0.26±0.12, *a* = 1.07±0.74, *r* = 0.4), estimated from HMM firing rate fits on all ongoing trials in each session (varying from 73 to 129). X-axis: ensemble size; Y-axis: dimensionality. C: Fraction of variance explained by each principal eigenvalue for the ensemble in panel A during evoked activity. Principal eigenvalues for the four tastes sucrose (S, orange), sodium chloride (N, yellow), citric acid (C, cyan), and quinine (Q, blue) are presented (corresponding to the color-coded dots in panel D). X-axis: eigenvalue number; Y-axis: percentage of variance explained by each eigenvalue. D: Dimensionality of neural activity across all ensembles in the empirical dataset during evoked activity (notations as in panel B, linear regression: *d = b·N +a, b* = 0.13 ± 0.03, *a* = 1.27 ± 0.19, *r* = 0.39), estimated from HMM firing rate fits on evoked trials in each condition (varying from 7 to 11 trials across sessions for each tastant). E: The slope of the linear regression of dimensionality (*d*) vs. ensemble size (*N*) as a function of the length of the trial interval and the number of trials used to estimate the dimensionality. X-axis: length of trial interval [s]; Y-axis: number of trials. F: Time course of the trial-matched slopes of *d* vs. *N*, evaluated with 200 ms bins in consecutive 1 s intervals during ongoing (black curve, *t* < 0) and evoked periods (red curve, *t* > 0; error bars represent SD). A significant time course is triggered by stimulus presentation (see Results for details). The slopes of the empirical dataset (thick curves) were smaller than the slope of the shuffled dataset (dashed curves). X-axis: time from stimulus onset at t = 0 [s]; Y-axis: slope of d vs. N. G: Distribution of pair-wise correlations in simultaneously recorded ensembles (black and red histograms for ongoing and evoked activity, respectively) and shuffled ensembles (brown and pink dashed histograms for ongoing and evoked activity, respectively) from 200 ms bins. X-axis: correlation; Y-axis: frequency. H: Distribution of pair-wise correlations from HMM states during ongoing (black) and evoked activity (red) for all simultaneously recorded pairs of neurons. X-axis: correlation; Y-axis: frequency.

### 3.3 Stimulus-induced reduction of dimensionality

We matched the number and duration of the trials for each data point and ran a two-way ANOVA with condition (ongoing vs. evoked) and ensemble size as factors. Both the main dimensionality (*F*_1,202_ = 11.93, *p* < 0.001) and the slope were significantly smaller during evoked activity (test of interaction, *F*_6,202_ = 5.09, *p* < 10^−4^). There was also a significant effect of ensemble size (*F*_6,202_ = 18.72, *p* < 10^−14^), confirming the results obtained with the separate regression analyses. These results suggest that stimuli induce a reduction of the effective space visited by the firing rate vector during evoked activity. This was confirmed by a paired sample analysis of the individual dimensionalities across all 27 × 4 = 108 ensembles (27 ensemble times 4 gustatory stimuli; *p* < 0.002, Wilcoxon signed-rank test).

### 3.4 Dimensionality is larger in ensembles of independent neurons

The dimensionality depends on the pair-wise correlations of simultaneously recorded neurons. Shuffling neurons across ensembles would destroy the correlations (beyond those expected by chance), and would give a measure of how different the dimensionality of our datasets would be compared to sets of independent neurons. We measured the dimensionality of surrogate datasets obtained by shuffling neurons across sessions; because shuffling destroys the structure of the hidden states, firing rates in bins of fixed duration (200 ms) were used to estimate the dimensionality (see Methods for details). As expected, the slope of *d* vs. *N* was larger in the shuffled datasets compared to the simultaneously recorded ensembles (not shown) during both ongoing activity (*b_shuff_* = 0.67 ± 0.06 vs. *b_data_* = 0.60 ± 0.01; mean ± SD, Mann-Whitney test, *p* < 0.001, 20 bootstraps), and evoked activity (*b_shuff_* = 0.36 ± 0.07 vs. **b_data_** = 0.29 ± 0.01; *p* < 0.001). Especially during ongoing activity, this result was accompanied by a narrower distribution of pair-wise correlations in the shuffled datasets compared to the simultaneously recorded datasets (Fig. 3G), and is consistent with an inverse relationship between dimensionality and pair-wise correlations (see e.g. Eq. (2.10)).

### 3.5 Time course of dimensionality as a function of ensemble size

Unlike ongoing activity, the dependence of dimensionality on ensemble size (the slope of the linear regression of *d* vs. *N*) was modulated during different epochs of the post-stimulus period (Fig. 3F, full lines; two-way ANOVA; main effect of time *F*_(4,495)_ = 3.80, *p* < 0.005; interaction time × condition: *F*_(4,495)_ = 4.76, *p* < 0.001). In particular, the dependence of *d* on the ensemble size *N* almost disappeared immediately after stimulus presentation in the simultaneously recorded, but not in the shuffled ensembles (trial-matched slope in the first evoked second: *b_evoked_* = 0.07 ± 0.01 vs *b_shuff_* = 0.19 ± 0.07) and converged to a stable value after approximately 1 second (slope after the first second *b_evoked_* = 0.38 ± 0.01; compare with a stable average slope during ongoing activity of *b_ongoing_* = 0.57 ± 0.01, Fig. 3F). Note that the dimensionality is larger when the firing rate is computed in bins (as in Fig. 3F) rather than in HMM states (as in Fig. 3B-D, where the slopes are about half than in Fig. 3F). The reason is that firing rates and correlations are approximately constant during the same HMM state, whereas they may change when estimated in bins of fixed duration that include transitions among hidden states. These changes tend to dilute the correlations resulting in higher dimensionality as predicted e.g. by Eq. (2.10). A comparison of the pair-wise correlations of binned firing rates (Fig. 3G) vs. those of firing rates in HMM states (Fig. 3H) confirmed this hypothesis. Also, if the argument above is correct, one would expect a dependence of dimensionality on (fixed) bin duration. We computed the correlations and dimensionality of binned firing rates for various bin durations and found that *r* increases and *d* decreases for increasing bin durations (not shown). However, the slope of *d* vs. *N* is always larger in ongoing than in evoked activity regardless of bin size (ranging from 10 ms to 5 s; not shown). This confirms the generality of the results of Fig. 3B-D, which were obtained using firing rate vectors in hidden states. To summarize our main results so far, we found that dimensionality depends on ensemble size during both ongoing and evoked activity, and such dependence is significantly reduced in the post-stimulus period. This suggests that while state sequences during ongoing activity explore a large portion of the available firing rate space, the presentation of a stimulus initially collapses the state sequence along a more stereotyped and lower-dimensional response [22, 24]. During both ongoing and evoked activity, the dimensionality is also different than expected by chance in a set of independent neurons (shuffled datasets).

**Figure 4.**
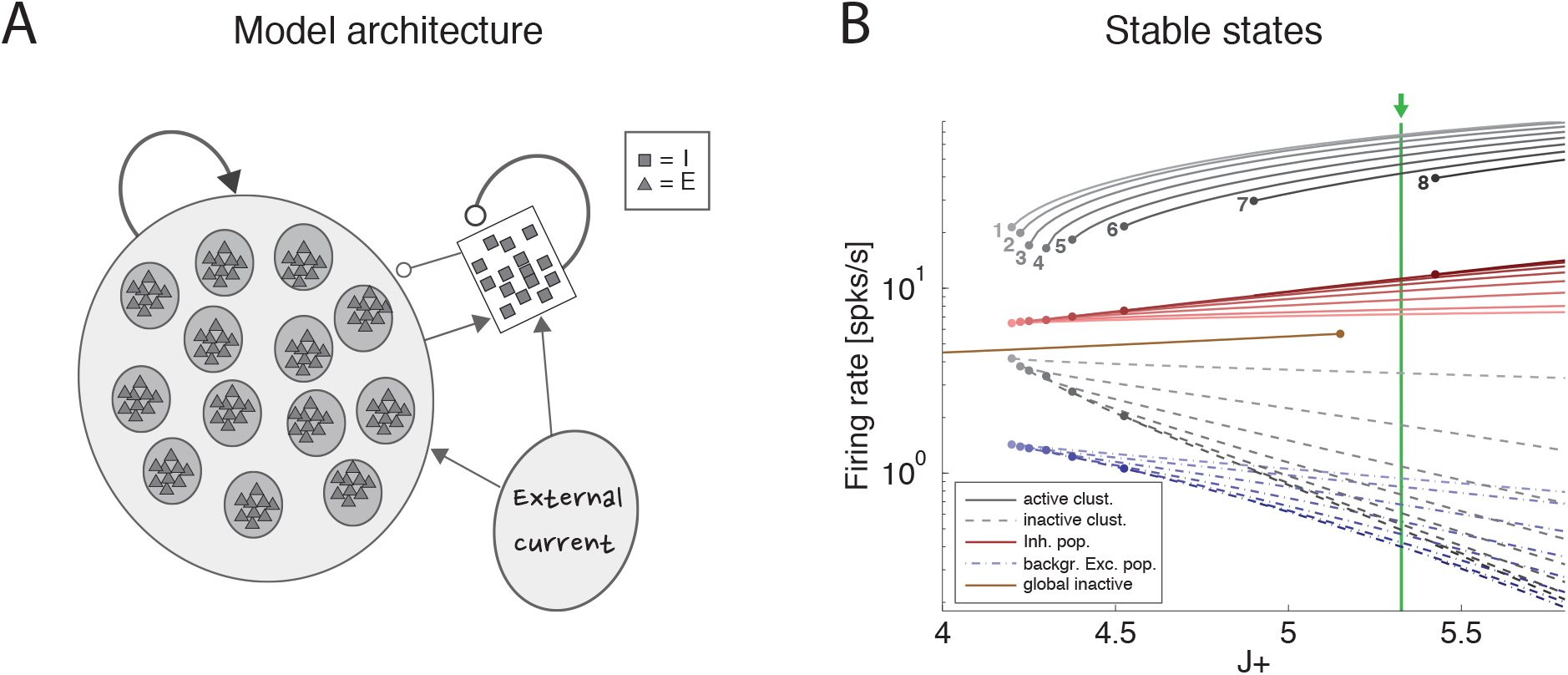
Recurrent network model. A: Schematic recurrent network architecture. Triangles and squares represent excitatory and inhibitory LIF neurons respectively. Darker disks indicate excitatory clusters with potentiated intra-cluster synaptic weights. B: Mean field solution of the recurrent network. Firing rates of the stable states for each subpopulation are shown as function of the intra-cluster synaptic potentiation parameter *J*_+_: firing rate activity in the active clusters (solid grey lines), firing rate in the inactive clusters (dashed grey lines), activity of the background excitatory population (dashed blue lines), activity of the inhibitory population (solid red lines). In each case, darker colors represent configurations with larger number of active clusters. Numbers denote the number of active clusters in each stable configuration. Configurations with 1 to 8 active clusters are stable in the limit of of infinite network size. A global configuration where all clusters are inactive (brown line) becomes unstable at the value *J*_+_ = 5.15. The vertical green line represents the value of *J*_+_ = 5.3 chosen for the simulations. X-axis: intra-cluster potentiation parameter *J*_+_ in units of *J_EE_*; Y-axis: Firing rate (spks/s).

### 3.6 Clustered spiking network model of dimensionality

To gain a mechanistic understanding of the different dimensionality of ongoing and evoked activity we have analyzed a spiking network model with clustered connectivity which has been shown to capture many essential features of the data [5]. In particular, the model reproduces the transitions among latent states in both ongoing and evoked activity. The network (see Section 2.8 for details) comprises *Q* clusters of excitatory neurons characterized by stronger synaptic connections within each cluster and weaker connections between neurons in different clusters. All neurons receive recurrent input from a pool of inhibitory neurons that keeps the network in a balanced regime of excitation and inhibition in the absence of external stimulation (Fig. 4A). In very large networks (technically, in networks with an infinite number of neurons), the stable configurations of the neural activity are characterized by a finite number of active clusters whose firing rates depend on the number clusters active at any given moment, as shown in Fig. 4B (where *Q* = 30). In a finite network, however, finite size effects ignite transitions among these configurations, inducing network states (firing rate vectors) on randomly chosen subsets of neurons that resemble the HMM states found in the data (Fig. 5; see [5] for details).

The dimensionality of the simulated sequences during ongoing and evoked activity was computed as done for the data, finding similar results. For the examples in Fig. 5, we found *d_ongoing_* = 4.0 for ongoing activity (Fig. 6A) between *d_evoked_* = 2.2 and *d_evoked_* = 3.2 across tastes during evoked activity (Fig. 6C). Across all simulated sessions, we found an average *d_ongoing_* = 2.9 ± 0.9 (mean±SD) for ongoing activity and *d_evoked_* = 2.4±0.7 for evoked activity. The model captured the essential properties of dimensionality observed in the data: the dimensionality did not differ across different tastes (one-way ANOVA, *p* > 0.2) and depended on ensemble size during both ongoing (Fig. 6B; slope = 0.36 ± 0.07, *r* = 0.77, *p* < 10^−4^) and evoked periods (Fig. 6D; slope = 0.12 ± 0.04, *r* = 0.29, *p* = 0.01). As for the data, the dependency on ensemble size was smaller for evoked compared to ongoing activity. We performed a trial-matched two-way ANOVA as done on the data and found, also in the model, a main effect of condition (ongoing vs. evoked: *F*_1,146_ = 22.1, *p* < 10^−5^), a main effect of ensemble size (*F*_6,146_ = 14.1, *p* < 10^−11^), and a significant interaction (*F*_6,146_ = 3.8, *p* = 0.001). These results were accompanied by patterns of correlations among the model neurons (Fig. 6E-F) very similar to those found in the data (Fig. 3G-H; see Section 3.10 for statistics of correlation values). As in the data, narrower distributions of correlations were found for binned firing rates (Fig. 6E) compared to firing rates in hidden states (Fig. 6F; compare with Figs. 3G-H, respectively). Moreover, shuffling neurons across datasets reduced the correlations (Fig. 6E, dashed), resulting in a larger slope of *d* vs. *N* (not shown). Finally, *d* during ongoing activity was always larger than during evoked activity also when computed on binned firing rates (not shown), as found in the data. Since the model was not fine-tuned to find these results, the different dimensionalities of ongoing and evoked activity, and their associated patterns of pair-wise correlations, are likely the consequence of the organization in clusters and of the ensuing dynamics during ongoing and evoked activity.

**Figure 5.**
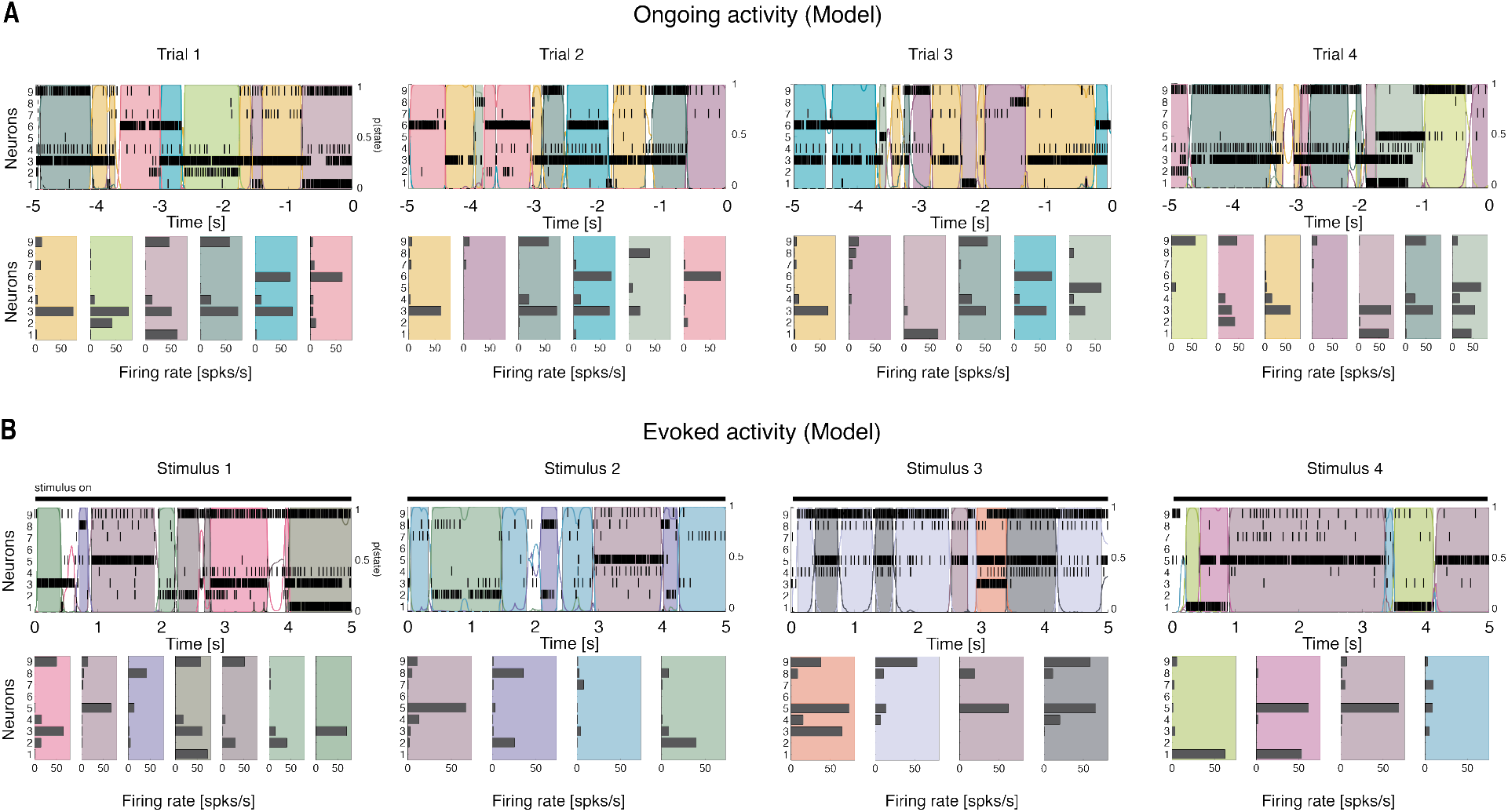
Ensemble activity in the recurrent network model is characterized by sequences of states. Representative trials from one ensemble of nine simultaneously recorded neurons sampled from the recurrent network, segmented according to their ensemble states (notations as in Fig. 1). A: Ongoing activity. B: Ensemble activity evoked by four different stimuli, modeled as an increase in the external current to selected clusters (the black line on top of the raster plot represents the “stimulus-on” period; see Methods for details).

### 3.7 Scaling of dimensionality with ensemble size and pair-wise correlations

The dependence of dimensionality on ensemble size observed in the data (Fig. 3B) and in the model (Fig. 6B) raises the question of whether or not the dimensionality would converge to an upper bound as one increases the number of simultaneously recorded neurons. In general, this question is important in a number of settings, related e.g. to coding in motor cortex [8, 14], performance in a discrimination task [13], or coding of visual stimuli [12]. We can attack this question aided by the model of Fig. 4, where we can study the effect of large numbers of neurons, but also the impact on dimensionality of a clustered network architecture compared to a homogeneous one, at parity of correlations and ensemble size.

**Figure 6.**
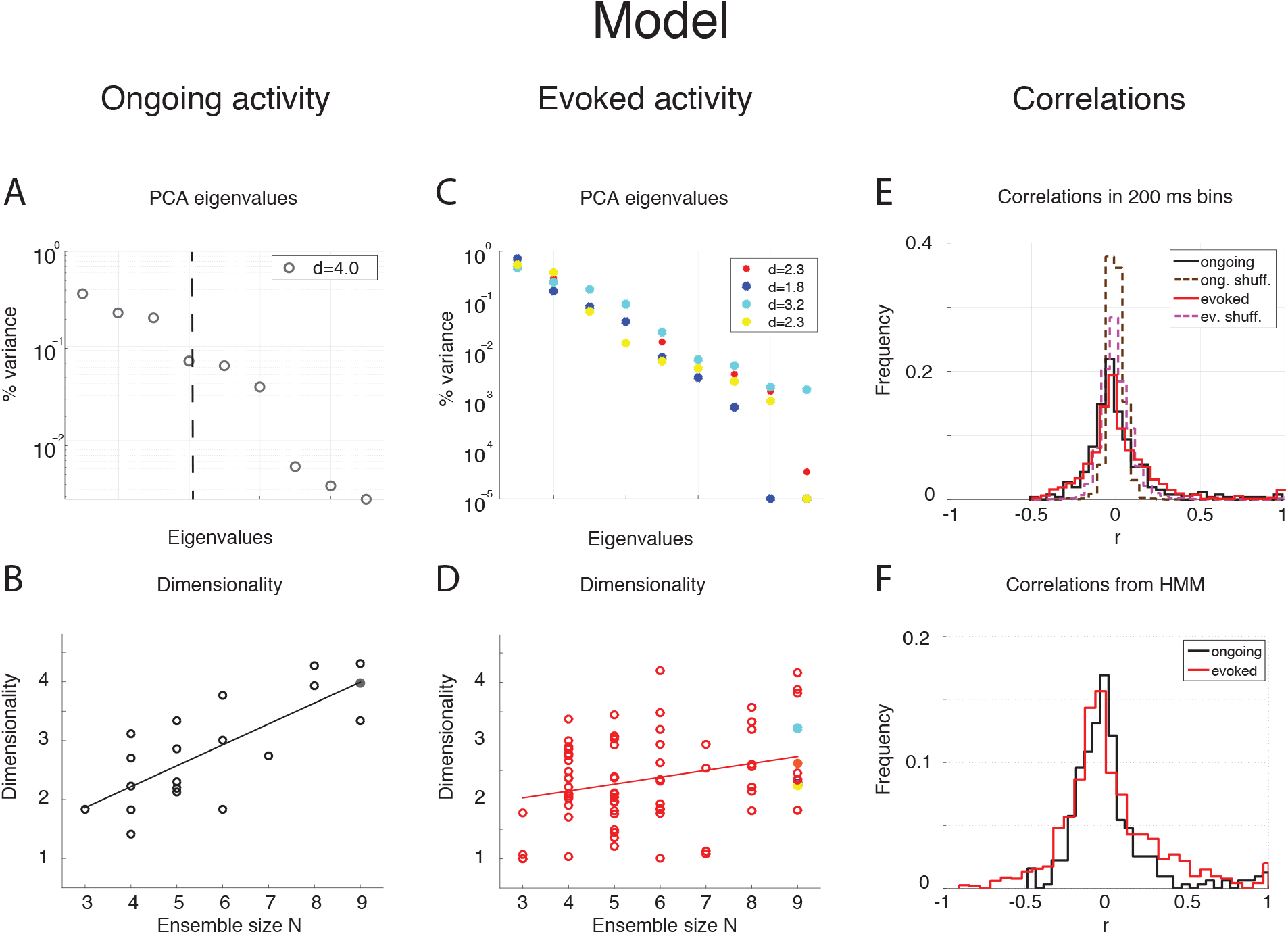
Dependence of dimensionality on ensemble size (model). A: Fraction of variance explained by each principal eigenvalue for an ensemble of 9 neurons during ongoing activity (corresponding to the filled dot in panel B) in the model network of Fig. 5 (notations as in Fig. 3A). B: Dimensionality of neural activity across all ensembles in the model during ongoing activity (linear regression fit, *d* = *b* · *N* + *a, b* = 0.36 ± 0.07, *a* = 0.80 ± 0.43, *r* = 0.77), estimated from HMM firing rate fits. X-axis: ensemble size; Y-axis: dimensionality. C: Fraction of variance explained by each principal eigenvalue for the ensemble in panel A during evoked activity. Principal eigenvalues for four stimuli are presented (corresponding to the color-coded dots in panel D). X-axis: eigenvalue number; Y-axis: percentage of variance explained by each eigenvalue. D: Dimensionality of neural activity across all ensembles in the model during evoked activity (notations as in panel B, linear regression: *d* = *b* · *N* + *a, b* = 0.12 ± 0.04, *a* = 1.70 ± 0.26, *r* = 0.29). E: Distribution of pair-wise correlations in simultaneously recorded ensembles from the clustered network model (black and red histograms for ongoing and evoked activity, respectively) and in shuffled ensembles (brown and pink dashed histograms for ongoing and evoked activity, respectively) from 200 ms bins. X-axis: correlation; Y-axis: frequency. F: Distribution of pair-wise correlations from HMM states during ongoing (black) and evoked activity (red) for all simultaneously recorded pairs of neurons. X-axis: correlation; Y-axis: frequency.

We consider first the case of a homogeneous network of neurons having no clusters and low pair-wise correlations, but having the same firing rates distributions (which were approximately log-normal, Fig. 7A) and the same mean pair-wise correlations as found in the data (*ρ* ∼ 0.01 – 0.2). This would require solving a homogeneous recurrent network self-consistently for the desired firing rates and correlations. As a proxy for this scenario, we generated 20 sessions of 40 Poisson spike trains having exactly the desired properties (including the case of independent neurons for which *ρ* = 0). Two examples with *ρ* = 0 and *ρ* = 0.1, respectively, are shown in Fig. 7B-C. Since in the asynchronous homogeneous network there are no transitions and hence no hidden states, the dimensionality was estimated based on the rate vectors in bins of 200 ms duration (using bin widths of 50 to 500 ms did not change the results; see Methods for details).

We found that the dimensionality grows linearly with ensemble size in the absence of correlations, but is a concave function of *N* in the presence of spike count correlations (circles in Fig. 7D). Thus, as expected, the presence of correlations reduces the dimensionality. A simple theoretical calculation mimicking this scenario shows that d in this case converges slowly to an upper bound that depends on the inverse of the square of the pair-wise correlations. For example, in the case of uniform correlations (*ρ*) and equal variances of the spike counts, Eq. (2.9) of Methods, 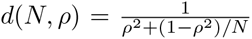, shows that *d = N* in the absence of correlations, and *d* ≤ 1/*ρ*^2^ for large networks in the presence of correlations. These properties remain approximately true in the case where the variances 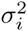 of the spike counts are drawn from a distribution with mean 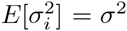 and variance 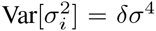. As Eq. (2.10) shows, in such a case dimensionality is reduced compared to the case of equal variances, for example 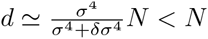 for large *N* when *ρ* = 0, *δρ* = 0.

The analytical results are shown in Fig. 7E (full lines correspond to Eq. (2.9)), together with their estimates (“+”) based on 1,000 data points (same number as trials in Fig. 7D; see Methods). The estimates are based on surrogate datasets with lognormal-distributed variances 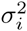 to mimic the empirical distribution of variances found in GC (not shown).

### 3.8 Estimation bias

Comparison of Fig. 7D with 7E shows that the dimensionality of the homogeneous network is underestimated compared to the theoretical value given by Eq. (2.9). This is due to a finite number of trials and the presence of unequal variances with spread *δσ*^4^ (“+” in Fig. 7E). As Fig. 7E shows, taking this into account will reduce the dimensionality to values comparable to those of the homogeneous network of Fig. 7D. The dimensionality in that case is well predicted by Eq. (2.17) (broken lines in Fig. 7E). The same Eq. (2.17) was fitted successfully to the data in Fig. 7D (dashed) by tuning 2 parameters to account for the unknown variance and correlation width of the firing rates (see Methods for details).

Empirically, estimates of the dimensionality Eq. (2.2) based on a finite number *N_T_* of trials tend to underestimate *d* (Fig. 2.4F). The approximate estimator Eq. (2.17) confirms that, for any ensemble size *N, d* is a monotonically increasing function of the number of trials (Fig. 8A). Note that this holds for the mean value of the estimator (Eq. (2.17)) over many datasets, not for single estimates, which could overestimate the true *d* (not shown). Eq. (2.17) also provides an excellent description of dimensionality as a function of firing rates variance *δσ*^4^ (Fig. 8B) and pair-wise correlations width *δρ*^2^ (Fig. 8C). In particular, the mean and the variance of the pair-wise correlations have an interchangeable effect on *d* (see Eq. (2.17)); they both decrease the dimensionality and so does the firing rate variance *δσ*^4^ (Fig. 8B).

### 3.9 Scaling of dimensionality in the presence of clusters

We next compared the dimensionality of the homogeneous networks activity to that predicted by the clustered network model of Fig. 4. To allow comparison with the homogeneous network, dimensionality was computed based on the spike counts in 200 ms bins rather than the HMMs firing rate vectors as in Fig. 6 (see Section 2.4 for details).

We found that the dependence of *d* on *N* in the clustered network depends on how the neurons are sampled. If the sampling is completely random, so that any neuron has the same probability of being added to the ensemble regardless of cluster membership, a concave dependence on *N* will appear, much like the case of the homogeneous network (Fig. 9A, dashed lines). However, if neurons are selected one from each cluster until all clusters have been sampled once, then one neuron from each cluster until all clusters have been sampled twice, and so on, until all the neurons in the network have been sampled, then the dependence of *d* on *N* shows an abrupt transition when *N = Q*, i.e., when the number of sampled neurons reaches the number of clusters in the network (Fig. 9A, full lines; see Fig 8B for raster plots with *Q* = 30 and *N* = 50). In the following, we refer to this sampling procedure as “ordered sampling”, as a reminder that neurons are selected randomly from each cluster, but the clusters are selected in serial order. For *N ≤ Q*, the dimensionality grows linearly with ensemble size in both ongoing (slope 0.24 ± 0.01, *r* = 0.79, *p* < 10^−10^, black line) and evoked periods (slope 0.19 ± 0.01, *r* = 0.84, *p* < 10^−10^; red line), and was larger during ongoing than evoked activity (trial-matched two-way ANOVA, main effect: *F*_1,948_ = 168, *p* < 10^−30^; interaction: *F*_5,948_ = 4.1, *p* < 0.001). These results are in keeping with the empirical and model results based on the HMM analysis (Fig. 3 and 6). However, in the case of ordered sampling, the dependence of dimensionality on ensemble size tends to disappear for *N ≥ Q* both during ongoing (slope 0.010 ± 0.003, *r* = 0.1, *p* < 0.001) and evoked periods (slope 0.009 ± 0.002, *r* = 0.13, *p* < 10^−4^; Fig. 9A, full lines). The average dimensionality over the range 30 ≤ *N* < 100 was significantly larger for ongoing, *d_ongoing_* = 8.74 ± 0.06, than for evoked activity, (trial-matched two-way ANOVA, main effect: *F*_1,2212_ = 488, *p* < 10^−30^), confirming that dimensionality is larger during ongoing than evoked activity also in this case. The difference in dimensionality between ongoing and evoked activity also holds in the case of random sampling on the entire range of *N* values (Fig. 9A, dashed lines), confirming the generality of this finding.

**Figure 7.**
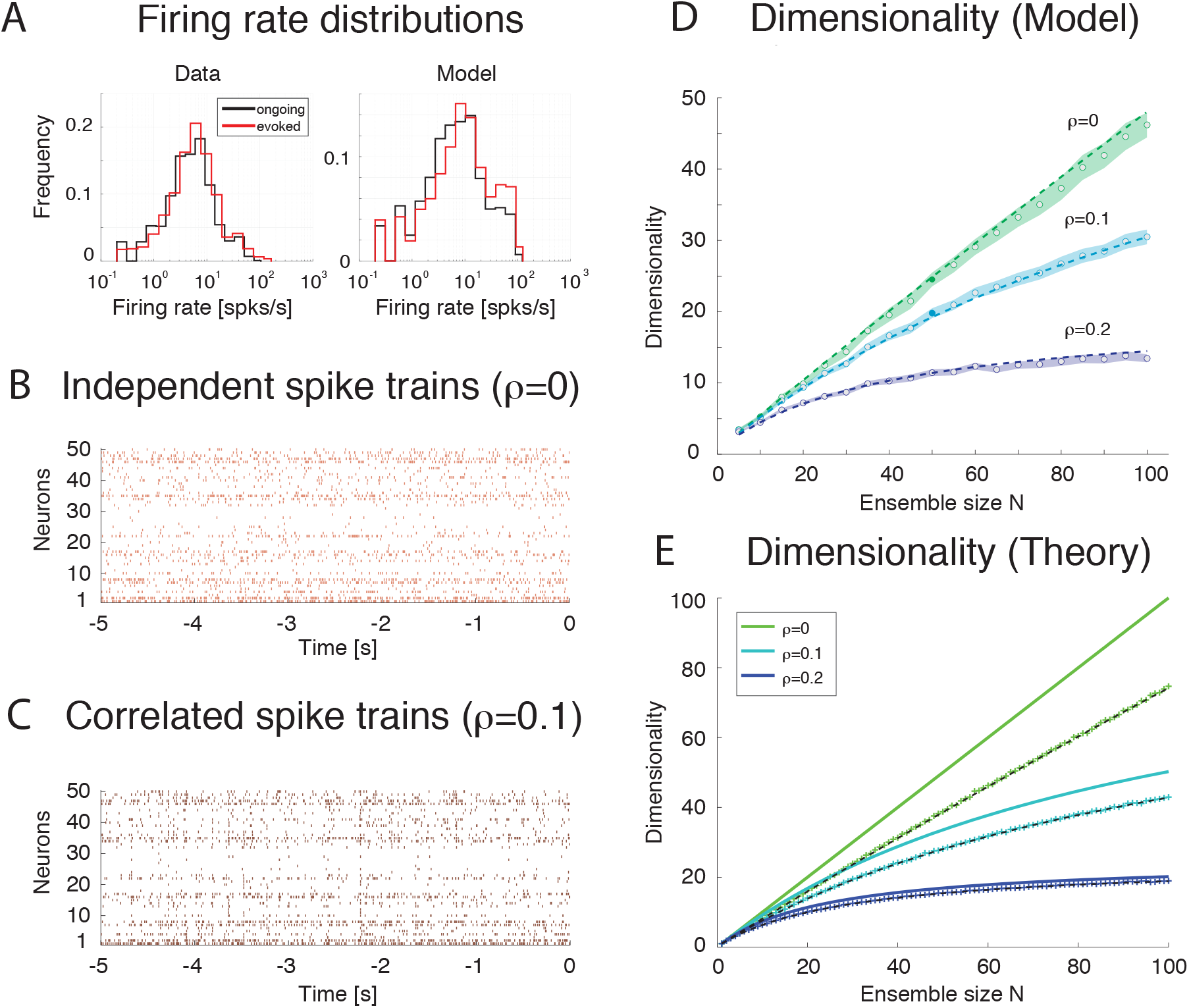
Dimensionality and correlation. A: Empirical single neuron firing rate distributions in the data (left) and in the model (right), for ongoing (black) and evoked activity (red). The distributions are approximately lognormal. X-axis: Firing rate (spks/s); Y-axis: density. B: Example of independent Poisson spike trains with firing rates matched to the firing rates obtained in simulations of the spiking network model. C: Example of correlated Poisson spike trains with firing rates matched to the firing rates obtained in simulations of the spiking network model. Pair-wise correlations of *ρ* = 0.1 were used (see Methods). X-axis: time [s]; Y-axis: neuron index. D: Dimensionality as a function of ensemble size N in an ensemble of Poisson spike trains with spike count correlations *ρ* = 0, 0.1, 0.2 and firing rates matched to the model simulations of Fig. 6. Dashed lines represent the fit of Eq. (2.17) to the data (with *δρ*^2^ = *αρ*^2^, σ^4^ = *δσ*^4^ = *β*), with best-fit parameters (mean±s.e.m.) *α* = 0.22 ± 10^−5^, *β* = 340 ± 8. Filled circles (from top to bottom): dimensionality of the data (raster plots) shown in panel B, C (shaded areas represent SD). X-axis: ensemble size; Y-axis, dimensionality. E: Theoretical prediction for the dependence of dimensionality on ensemble size N and spike count correlation *ρ* for the case of uniform correlation, Eq. (2.9) (thick lines; green to cyan to blue shades represent increasing correlations). “+” are dimensionality estimates from *N_T_* = 1,000 trials for each *N* (same *N_T_* as in panel D, each trial providing a firing rate value sampled from a log-normal distribution), in the case of log-normally distributed firing rate variances 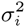 with mean *σ*^2^ = 40 (spk/s)^2^ and standard deviation 0.5*σ*^2^. Theoretical predictions from Eq. (2.17) match the estimated values in all cases (dashed black lines). X-axis: ensemble size N; Y-axis: dimensionality.

**Figure 8.**
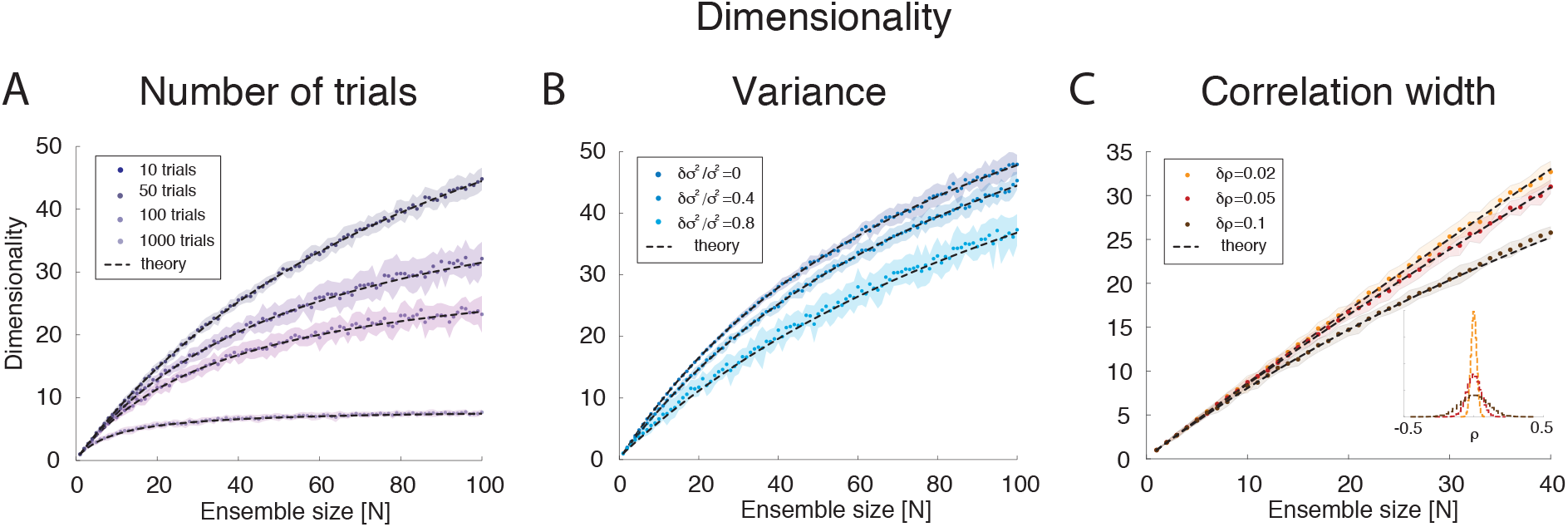
Dimensionality estimation. A: Dependence of dimensionality on the number of trials for variable ensemble size *N*, for fixed correlations *ρ* = 0.1 and firing rates variances 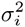 with mean *σ*^2^ and standard deviation *δσ*^2^ = 0.4*σ*^2^. Dashed lines: theoretical prediction, Eq. (2.17); dots: mean values from simulations of 20 surrogate datasets containing 10 to 1000 trials each (shaded areas: SD), with darker shades representing increasing number of trials. X-axis: ensemble size; Y-axis, dimensionality. B: Dependence of dimensionality on the spread of the firing rates variances for fixed correlations *ρ* = 0.1 and firing rate variance with mean *σ*^2^. Dashed lines: theoretical prediction, Eq. (2.17); dots: mean values from simulations of 20 surrogate datasets containing 1000 trial each (shaded areas: SD), with lighter shades representing increasing values of *δσ*^2^/*σ*^2^). X-axis: ensemble size; Y-axis, dimensionality. C: Dependence of dimensionality on the width 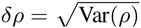 of pair-wise firing rate correlations (with zero mean), for firing rates variances 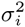 with mean *σ*^2^ and standard deviation *δσ*^2^ = 0.4*σ*^2^. Dashed lines: theoretical prediction, Eq. (2.17); dots: mean values from simulations of 20 surrogate datasets containing 1000 trials each (shaded areas: SD), with darker shades representing increasing values of *δρ*. Inset: distribution of correlation coefficients used in the main figure. X-axis: ensemble size; Y-axis, dimensionality. In all panels, *σ*^2^ = 40 (spk/s)^2^.

### 3.10 Dimensionality is larger in the presence of clusters

Intuitively, the dimensionality saturates at *N = Q* in the clustered network because additional neurons will be highly correlated with already sampled ones. For *N ≤ Q*, each new neurons activity adds an independent degree of freedom to the neural dynamics and thus increases its dimensionality. For *Q > N*, additional neurons are highly correlated with an existing neuron, adding little or no additional contribution to *d*. Indeed, compared to the low overall correlations found across all neuron pairs in the data (and used as *desiderata* for the homogeneous network), neurons belonging to the same model cluster had a much higher spike count correlation of *ρ* = 0.92 [0.56,0.96] (median and [25, 75]-percentile), while neurons belonging to different clusters had a much lower correlation of *ρ* ⋍ 0 [-0.10,0.06]. A negligible median correlation was typical: for example, negligible was the overall median correlation regardless of cluster membership (*ρ* ⋍ 0 [-0.109,0.083]); and the empirical correlation both during ongoing ([-0.047,0.051], with rare maximal values of *ρ* ∼ 0.5), and evoked activity ([-0.085,0.113], with rare maximal values of *ρ* ∼ 0.9). While we note the qualitative agreement of model and empirical correlations, we emphasize that these numbers were obtained using 200 ms bins and that they were quite sensitive to bin duration. In particular, the maximal correlations (regardless of sign) were substantially reduced for smaller bin durations (not shown). Plugging these values into a correlation matrix reflecting the clustered architecture and the sampling procedure used in Fig. 9B, we obtained the matrix shown in Fig. 9C, where pairwise correlations depend on whether or not the neurons belong to the same cluster (for the first 40 neurons, adjacent pairs belong to the same cluster; the last 10 neurons belong to the remaining clusters). It is natural to interpret such correlation matrix as the noisy observation of a block-diagonal matrix such that neurons in the same cluster have uniform correlation while neurons from different clusters are uncorrelated. For such a correlation matrix the dimensionality can be evaluated exactly (see Eq. (2.13)). In the approximation where all neurons have the same variance, this reduces to Eq. (2.14), i.e.

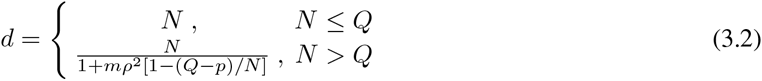

**Figure 9.**
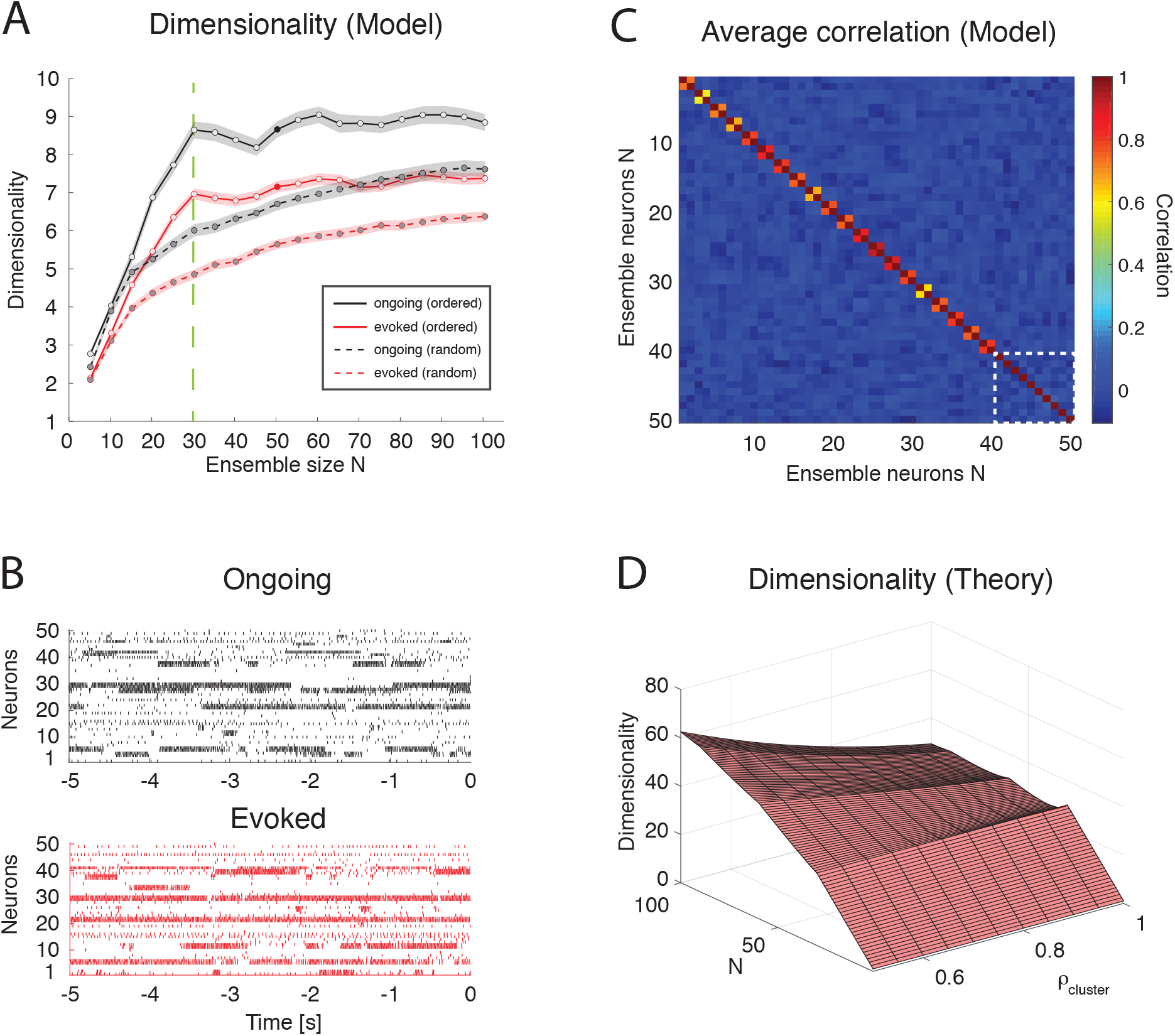
Dimensionality in a clustered network. A: Trial-matched dimensionality as a function of ensemble size in the recurrent network model (ongoing and evoked activity in black and red, respectively, with shaded areas representing s.e.m.). Filled lines represent ordered sampling, where ensembles to the left of the green vertical line (*N* = *Q* = 30) contain at most one neuron per cluster, while to the right of the line they contain one or more neurons from all clusters (filled circles indicate representative trials in panel B). Dashed lines represent random sampling of neurons, regardless of cluster membership. X-axis: ensemble size; Y-axis, dimensionality. B: Representative trial of an ensemble of 50 neurons sampled from the recurrent network in Fig. 4 during ongoing activity (upper plot, in black) or evoked activity (lower plot, in red) for the case of ordered sampling (full lines). Neurons are sorted according to their cluster membership (adjacent neuron pairs with similar activity belong to the same cluster, for neurons #1 up to #40; the last ten neurons are sampled from the remaining clusters). X-axis: time to stimulus presentation at *t* = 0 (s); Y-axis: neuron index. C: Average correlation matrix for twenty ensembles of *N* = 50 neurons from the clustered network model with *Q* = 30. For the first 40 neurons, adjacent pairs belong to the same cluster; the last 10 neurons (delimited by a dashed white square) belong to the remaining clusters (neurons are ordered as in panel B). Thus, neurons 1, 3, 5,… 39 (20 neurons) belong to the first 20 clusters; neurons 2, 4, 6,… 40 (20 neurons) belong also the first 20 clusters; and neurons 41, 42, 43,…, 50 (10 neurons) belong to the remaining 10 clusters. X-axis, Y-axis: neuron index. D: Plot of Eq. 2.13 giving *d* vs. *N* and *ρ* (uniform within-cluster correlations) for the sampling procedure of panel B. X-axis: ensemble size *N*; Y-axis: dimensionality.

where *N* = *mQ* + *p*. This formula is plotted in Fig. 9D for relevant values of *ρ* and *N* and it explains the origin of the abrupt transition in dimensionality at *Q = N*. (The reasons for a dimensionality lower than *Q* for *N ≤ Q* in the data see Fig. 9A are, also in this case, the finite number of data points (250) used for its estimation and the non-uniform distributions of firing rate variances and correlations).

Note that the formula also predicts cusps in dimensionality (which become local maxima for large *ρ*) whenever the ensemble size is an exact multiple of the number of clusters. This is also visible in the simulated data of Fig. 9A, where local maxima seem to appear at *N* = 30,60,90 with *Q* = 30 clusters. It is also worth mentioning that for low intra-cluster correlations the dependence on *N* predicted by Eq. (2.14) becomes smoother and the cusps harder to detect (not shown), suggesting that the behavior of a clustered network with weak clusters tends to converge to the behavior of a homogeneous asynchronous network – therefore lacking sequences of hidden states. Thus, the complexity of the network dynamics is reflected in how its dimensionality scales with *N*, assuming that one may sample one neuron per cluster (i.e., via “ordered sampling”). Even though it is not clear how to perform ordered sampling empirically (see Discussion), this result is nevertheless useful since it represents an upper bound also in the case of random sampling (see Fig. 9A, dashed lines). Eq. (2.14) predicts that *d ≤ Q/ρ*^2^, with this value reached asymptotically for large *N*. In the case of random sampling, growth to this bound is even slower (Fig. 9A). For comparison, in a homogeneous network *d* ≤ 1/*ρ*^2^ from Eq. (2.9), a smaller bound by a factor of *Q*. Finally, homogeneous dimensionality is dominated by clustered dimensionality also in the more realistic case of non-uniform variances, where similar bounds are found in both cases (see Section 2.4 for details).

## 4 Discussion

In this paper we have investigated the dimensionality of the neural activity in the gustatory cortex of alert rats. Dimensionality was defined as a collective property of ensembles of simultaneously recorded neurons that reflects the effective space occupied by the ensemble activity during either ongoing or evoked activity. If one represents ensemble activity in terms of firing rate vectors, whose dimension is the number of ensemble neurons *N*, then the collection of rate vectors across trials takes the form of a set of points in the *N*-dimensional space of firing rates. Roughly, dimensionality is the minimal number of dimensions necessary to provide an accurate description of such set of points, which may be localized on a lower-dimensional subspace inside the whole firing rate space.

One of the main results of this paper is that the dimensionality of evoked activity is smaller than that of ongoing activity, i.e., stimulus presentation quenches dimensionality. More specifically, the dimensionality is linearly related to the ensemble size, with a significantly larger slope during ongoing activity compared to evoked activity (compare Fig. 3B and 3D). We explained this phenomenon using a biologically plausible, mechanistic spiking network model based on recurrent connectivity with clustered architecture. The model was recently introduced in [5] to account for the observed dynamics of ensembles of GC neurons as sequences of metastable states, where each state is defined as a vector of firing rates across simultaneously recorded neurons. The model captures the reduction in trial-to-trial variability and the multiple firing rates attained by single neurons across different states observed in GC upon stimulus presentation. Here, the same model was found to capture also the stimulus-induced reduction of dimensionality. While the set of active clusters during ongoing activity varies randomly, allowing the ensemble dynamics to explore a large portion of firing rate space, the evoked set of active clusters is limited mostly to the stimulus-selective clusters only (see [5] for a detailed analysis). The dynamics of cluster activation in the model thus explains the more pronounced dependence of dimensionality on ensemble size found during ongoing compared to evoked activity.

We presented a simple theory of how dimensionality depends on the number of simultaneously recorded neurons *N*, their firing rate correlations, their variance, and the number and duration of recording trials. We found that dimensionality increases with *N* and decreases with the amount of pair-wise correlations among the neurons (e.g., Fig. 8C). At parity of correlations, dimensionality is maximal when all neurons have the same firing rate variance, and it decreases as the distribution of count variances becomes more heterogeneous (e.g., Fig. 8B). The estimation of dimensionality based on a finite dataset is an increasing function of the number of trials (Fig. 8A). Finally, introducing clustered correlations in the theory, and sampling one neuron per cluster as in Fig. 9B, results in cusps at values of *N* that are multiples of the number of clusters (Fig. 9D), in agreement with the predictions of the spiking network model (Fig. 9A, full lines).

### 4.1 Dimensionality scaling with ensemble size

The increased dimensionality with sample size, especially during ongoing activity, was found empirically in datasets with 3 to 9 neurons per ensemble, but could be extrapolated for larger *N* in a spiking network model with homogeneous or clustered architecture. In homogeneous networks with finite correlations the dimensionality is predicted to increase sub-linearly with *N* (Eq. 2.9), whereas in the clustered network it may exhibit cusps at multiple values of the number of clusters (Fig. 9A), and would saturate quickly to a value that depends on the ratio of the number of clusters *Q* and the amount of pair-wise correlations, *d ≤ Q/ρ*^2^. Testing this prediction requires the ability to sample neurons one from each cluster, until all clusters are sampled, and seems beyond the current recording techniques. However, looking for natural groupings of neurons based on response similarities could uncover spatial segregation of clusters [53] and could perhaps allow sampling neurons according to this procedure. Moreover, the model predicts a slower approach to a similar bound also in the case of random sampling.

Dimensionality in a homogeneous network is instead bounded by 1/*ρ*^2^, and hence it is a factor *Q* smaller than in the clustered network. Dimensionality is maximal in a population of independent neurons (*ρ* = 0), where it grows linearly with *N*; however, neurons of recurrent networks have wide-ranging correlations (see e.g. Fig. 6E-F and its empirical counterpart, Fig. 3G-H). Since the presence of even low correlations can dramatically reduce the dimensionality (see e.g. Fig. 7D), the neural activity in a clustered architecture can reach much higher values at parity of correlations, representing an intermediate case between a homogeneous network and a population of independent neurons.

Evidence for the presence of spatial clusters has been recently been reported in the prefrontal cortex based on correlations analyses [53]. An alternative possibility is that neural clusters are not spatially but functionally arranged, and cluster memberships vary with time and task complexity [54]. Can our model provide indirect tools to help uncover the presence of clusters? A closer look at Figs. 6E-F reveal a small peak at large correlations due to the contribution of highly correlated neurons belonging to the same cluster. This peak would be absent in a homogenous network and thus is the signature of a clustered architecture. However, such peak is populated by only small fraction (1/*Q*) of the total number of neuron pairs, which hinders its empirical detection (no peak at large correlations is clearly visible in our data, see Fig. 3G-H).

### 4.2 Dimensionality and trial-to-trial variability

Cortical recordings from alert animals show that neurons produce irregular spike trains with variable spike counts across trials [21, 55, 56]. Despite many efforts, it remains a key issue to establish whether variability is detrimental [57, 58] or useful [59] for neural computation.

Trial-to-trial variability is reduced during preparatory activity [60], during the presentation of a stimulus [61], or when stimuli are expected [17], a phenomenon that would not occur in a population of independent or homogeneously connected neurons [62]. Recent work has shown that the stimulus-induced reduction of trial-to-trial variability can be due to spike-frequency adaptation in balanced networks [63] or to slow dynamic fluctuations generated in a recurrent spiking networks with clustered connectivity [5, 62, 64]. In clustered network models, slow fluctuations in firing rates across neurons can ignite metastable sequences of neural activity, closely resembling metastable sequences observed experimentally [1–7]. The slow, metastable dynamics of cluster activation produces high variability in the spike count during ongoing activity. While cluster activations occur at random times during ongoing activity periods, stimulus presentation locks cluster activation at its onset, leading to a decrease in trial-to-trial variability.

Similarly, a stimulus-induced reduction of dimensionality is obtained in the same model. In this case, preferred cluster activation due to stimulus onset generates an increase in pair-wise correlations that reduce dimensionality. Note that the two properties (trial-to-trial variability and dimensionality) are conceptually distinct. An ensemble of Poisson spike trains can be highly correlated (hence have low dimensionality), yet the Fano Factor of each spike train will still be 1 (hence high), independently of the correlations among neurons. In a recurrent network, however, dimensionality and trial-to-trial variability may become intertwined and exhibit similar properties, such as the stimulus-induced reduction observed in a model with clustered connectivity. A deeper investigation of the link between dimensionality and trial-to-trial variability in recurrent networks is left for future studies.

### 4.3 Alternative definitions of dimensionality

Following [10] we have defined dimensionality (Eq. (2.2)) as the dimension of an effective linear subspace of firing rate vectors containing the most variance of the neural activity. It differs from the typical dimensionality reduction based on PCA in that the latter retains only the number of eigenvectors explaining a predefined amount of variance (see e.g. [65, 66]), because Eq. (2.2) includes contribution from all eigenvalues. Moreover, we have computed the firing rate correlations in bins of variable width that match the duration of the HMM states. Although our main results do not depend on bin size (see Section 3.5), the actual value of dimensionality decreases with increasing bin duration. Thus, any choice of bin size (e.g., 200 ms in Fig. 3F-G) remains somewhat arbitrary. A better method is to use a variable bin size as dictated by the HMM analysis, as done in Fig. 3B-D. This method also prevents diluting correlations among firing rates that would occur if one neuron were to change state inside the current bin, because during a hidden state the firing rates of the neurons are constant (by definition). Thus, this provides a principled adaptive procedure for selecting the bin size and eliminates the dependence of dimensionality on the bin width used for the analysis.

Other definitions of neural dimensionality have been proposed in the literature, which aim at capturing different properties of the neural activity, typically during stimulus-evoked activity. A measure of dimensionality related to ours, and referred to as “complexity,” was introduced in [12]. According to their definition, population firing rate vectors from all evoked conditions were first decomposed along their kernel Principal Components [67]. A linear classifier was then trained on an increasing number of leading PCs in order to perform a discrimination task, where the number of PCs used was defined as the complexity of the representation. In general, the classification accuracy improves with increasing complexity, and it may saturate when all PCs containing relevant features are used – with the remaining PCs representing noise or information irrelevant to the task. Reaching high accuracy at low complexity implies good generalization performance, i.e., the ability to classify novel variations of a stimulus in the correct category. Neural representations in monkey inferotemporal cortex (IT) were found to require lower complexity than in area V4, confirming ITs premier role in classifying visual objects despite large variations in shape, orientation and background [12]. Complexity relies on a supervised algorithm and is an efficient tool to capture the generalization properties of evoked representations (see e.g. [68]) for its relevance to visual object recognition).

A second definition of dimensionality, sometimes referred to as “shattering dimensionality” in the Machine Learning literature, was used to assess the discrimination properties of the neural representation [13]. Given a set of *p* firing rate vectors, one can split them into two classes (e.g. white and black colorings) in 2^*p*^ different ways, and train a classifier to learn as many of those binary classification labels as possible. The shattering dimensionality is then defined as (the logarithm of) the largest number of binary classifications that can be implemented. This measure of dimensionality was found to drop significantly in monkey prefrontal cortex during error trials in a recall task and thus predicts the ability of the monkey to correctly perform the task [13].

A flexible and informative neural representation is one that achieves a large shattering dimensionality (good discrimination) while keeping a low complexity (good generalization). Note that both complexity and shattering dimensionality represent measures of classification performance in task-related paradigms, and their definition requires a set of evoked conditions to be classified via a supervised learning algorithm. While both definitions could be applied to neural activity in our stimulus-evoked data, their interpretation cannot be extended to periods of ongoing activity, as the latter is not associated to desired targets in a way that can be learned by a classification algorithm. Since our main aim was to compare the dimensionality of ongoing and evoked activity, the unsupervised approach of [10] and their notion of “effective” dimensionality was better suited for our analysis. A related definition of dimensionality has been used by [14] to investigate neural representations of movements in motor cortex.

Many measures of dimensionality used in the literature (including ours and some of those discussed above) are based on pair-wise correlations. However, neural activity is known to give rise also to higher-order correlations [69]. Given that the extent and relevance of higher-order correlations is actively debated [70, 71], it would be useful to include them in measures of dimensionality. This is left for a future study.

### 4.4 Ongoing activity and task complexity

The relationship between ongoing and stimulus-evoked activity has been linked to the functional connectivity of local cortical circuits, and their mutual relationship has been the object of both theoretical and experimental investigations, often with contrasting conclusions (e.g., [5, 72–77]. Here, we have focused on the dimensionality of ongoing and evoked activity and have shown that neural activity during ongoing periods occupies a space of larger dimensionality compared to evoked activity. Although based on a different measure of dimensionality, recent results on the relation between the dimensionality of evoked activity and task complexity suggest that evoked dimensionality is roughly equal to the number of task conditions [13]. It is natural to ask whether the dimensionality of ongoing activity provides an estimate of the complexity of the hardest task that can be supported by the neural activity. Moreover, based on the clustered network model, the presence of clusters imposes an upper value *d ≤ Q/ρ*^2^ during ongoing activity, suggesting that a discrimination task with up to ∼ *Q* different conditions may be supported. The experience of taste consumption is by itself multidimensional, including both chemo- and oro-sensory aspects (i.e., taste identity [24] and concentration [78], texture, temperature [79, 80], taste and odor mixtures [81]) and psychological aspects (hedonic value [22, 24, 82]), anticipation [17], novelty [83] and satiety effects [84]. It is tempting to speculate that neural activity during ongoing periods explores these different dimensions, while evoked activity is confined to the features of the particular taste being delivered in a specific context. Establishing a precise experimental and theoretical link between the number of clusters and task complexity is an important question left for future studies.

## Acknowledgments

This work was supported by a National Institute of Deafness and Other Communication Disorders Grant K25-DC013557 (LM), by the Swartz Foundation Award 66438 (LM), by National Institute of Deafness and Other Communication Disorders Grant R01-DC010389 (AF), by a Klingenstein Foundation Fellowship (AF), and partly by a National Science Foundation Grant IIS1161852 (GLC). We thank Dr. Stefano Fusi and Memming Park for useful discussions and David Ecker at the Research Technologies DoIT of Stony Brook University for access to its computational resources.

